# Single-cell transcriptomics of allo-reactive CD4^+^ T cells over time reveals divergent fates during gut GVHD

**DOI:** 10.1101/2020.03.08.978239

**Authors:** Jessica A. Engel, Hyun Jae Lee, Cameron G. Williams, Rachel Kuns, Stuart Olver, Lianne I. M. Lansink, Megan S. F. Soon, Stacey B. Andersen, Joseph E. Powell, Valentine Svensson, Sarah A. Teichmann, Geoffrey R Hill, Antiopi Varelias, Motoko Koyama, Ashraful Haque

## Abstract

Acute gastrointestinal Graft-versus-Host-Disease (GVHD) is a primary determinant of mortality after allogeneic hematopoietic stem-cell transplantation (alloSCT). It is mediated by alloreactive donor CD4^+^ T cells that differentiate into pathogenic subsets expressing IFNγ, IL-17A or GM-CSF, and is regulated by subsets expressing IL-10 and/or Foxp3. Developmental relationships between T-helper states during priming in mesenteric lymph nodes (mLN) and effector function in the GI tract remain undefined at genome-scale. We used scRNA-seq and computational modelling to create an atlas of putative differentiation pathways during GVHD. Computational trajectory inference suggested emergence of pathogenic and regulatory states along a single developmental trajectory in mLN. Importantly, we identified an unexpected second trajectory, categorised by little proliferation or cytokine expression, reduced glycolysis, and high TCF1 expression. TCF1^hi^ cells upregulated α4β7 prior to gut migration and failed to express cytokines therein. Nevertheless, they demonstrated recall potential and plasticity following secondary transplantation, including cytokine or Foxp3 expression, but reduced TCF1. Thus, scRNA-seq revealed divergence of allo-reactive CD4^+^ T cells into quiescent and effector states during gut GVHD, reflecting putative heterogenous priming *in vivo*. These findings, the first at a single-cell level during GVHD over time, can now be used to interrogate T cell differentiation in patients undergoing alloSCT.

## Introduction

Allogeneic hematopoietic stem cell transplantation (alloSCT) is a curative therapy for a range of leukemias, due to the capacity of donor T cells within the transplant to kill tumour cells – known as the graft-versus-leukaemia (GVL) effect. Unfortunately, donor T cells can also damage non-cancerous host tissue, particularly the gastrointestinal (GI) tract, liver and skin, causing the serious condition, graft-versus-host-disease (GVHD). Acute GVHD in the GI tract remains the primary determinant of GVHD severity and risk of death (1). Thus, a primary goal for alloSCT is the prevention of acute gut GVHD while preserving GVL. We previously showed in pre-clinical models that donor CD4^+^ T cells are initially activated by recipient non-hematopoietic antigen-presenting cells (APC) within the gut, including epithelial cells that upregulate MHCII molecules (2, 3). Following this, donor-derived colonic dendritic cells (DC) also prime donor CD4^+^ T cells in mesenteric lymph nodes (mLN) and trigger T helper-cell differentiation, which serves to amplify and exacerbate GVHD (4). Using TCR transgenic T cells specific for a single allo-peptide, TEa cells (5), we revealed that donor CD4^+^ T cells within the same alloSCT recipient differentiate into multiple cellular states that express pro-inflammatory and pathogenic Th1/Th17-associated cytokines, including IFNγ and IL-17A, or the master transcription factor for induced regulatory T (iTreg) cells, Foxp3 (4). While cytokines such as IL-6 and IL-12 control differentiation of donor CD4^+^ T cells, fate-mapping studies based on the *IL-17a* promoter suggested complex dynamic relationships between apparent helper subsets (6). Thus, differentiation of allo-reactive donor CD4^+^ T cells is characterised by complexity, both in terms of dynamics and multiple cellular states adopted, neither of which have been explored at genome-scale.

Single-cell mRNA sequencing (scRNA-seq) enables unbiased genome-wide assessment of individual T cells without reliance upon pre-determined protein markers or genes. ScRNA-seq was previously employed to examine heterogeneity in CD4^+^ T cells isolated from *IL-17a* reporter mice undergoing experimental autoimmune encephalomyelitis (EAE) (7). Subsequently, scRNA-seq was used to examine CD4^+^ T cell differentiation during house dust mite-induced allergy (8), protozoan parasite infection (9), as well as CD8^+^ T cells in viral infections and cancer (10–12). Many of these studies were cross-sectional, offering insight into heterogeneity amongst clonal TCR transgenic cells at a single timepoint. We previously examined CD4^+^ T cell transcriptomes over a range of time-points during experimental malaria, and employed computational approaches to re-construct the dynamics of Th1 versus Tfh differentiation. Using an approach based on Bayesian Gaussian Process Latent Variable Modelling (bGPLVM), we identified a bifurcation point between two trajectories, and revealed a role for T cell extrinsic factors in governing Th1/Tfh fate (9). More recently, we employed scRNA-seq to reveal heterogeneity and tissue adaptation of thymic Tregs and colonic CD4^+^ T cells during steady-state in mice and humans (13, 14). Here, we examined donor antigen presenting cell (APC)-mediated differentiation of allo-reactive donor CD4^+^ T cells in acute gut GVHD using droplet-based scRNA-seq and computational modelling.

## Results

### Allo-reactive donor CD4^+^ T cells acquire heterogeneous pro-inflammatory, regulatory and uncharacterised states during acute GVHD

We previously established a pre-clinical model of acute GVHD (4), in which the donor CD4^+^ T cell responses were targeted to a single, host-derived, allo-reactive peptide presented by donor MHCII molecules - TEa TCR transgenic CD4^+^ T cells (B6 background) exhibit specificity for a BALB/c-derived Ea peptide from the MHCII molecule, I-E^d^, when presented by the donor (B6) MHCII molecule, I-A^b^ (Figure 1A). Here, we exposed BALB/c mice to total body irradiation and provided an MHC-mismatched B6 bone marrow transplant (containing donor T cells). 12-days later, once donor allo-presenting APCs had developed (4), TEa cells were transferred. TEa cell priming occurred specifically within the mesenteric lymph nodes (mLN), and triggered differentiation, as expected by day 4, into subsets with a strong capacity to express pro-inflammatory cytokines IFNγ or IL-17A upon re-stimulation, or in rarer cases (~1%) express Foxp3 (Figure 1A & B). We also examined the frequency of TEa cells expressing IFNγ, IL-17A or Foxp3 directly *ex vivo*, without further stimulation (Figure 1C). Although some TEa cells expressed IFNγ, IL-17A or Foxp3 directly *ex vivo* (Figure 1C), the majority did not at this early time-point in mLN. Therefore, although T helper cell differentiation had occurred, use of three canonical T helper markers was insufficient for characterising the fate of most TEa cells in mLN during acute gut GVHD.

**Figure 1.**
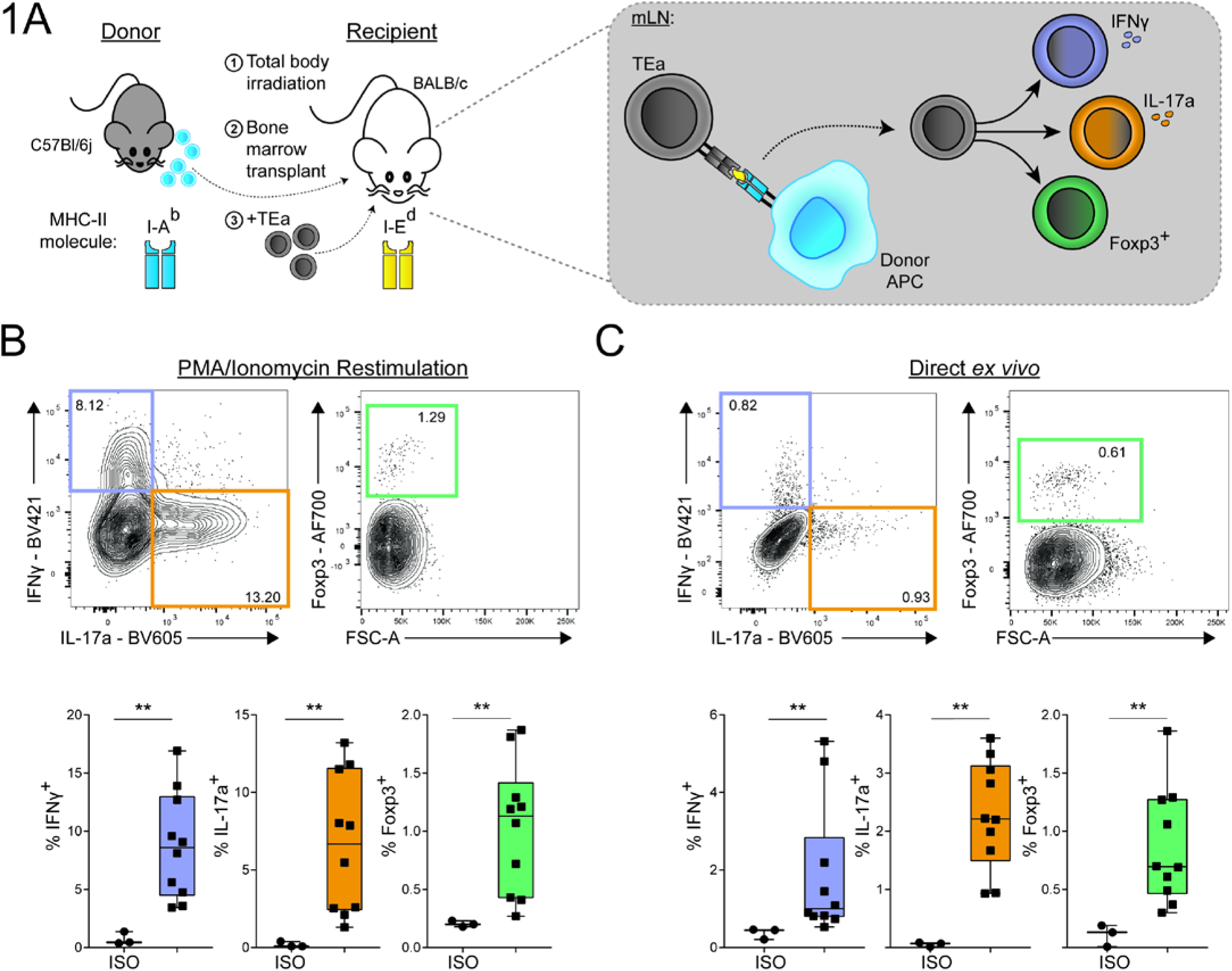
Allo-reactive donor CD4^+^ T cell clones acquire pro-inflammatory and regulatory states during acute GVHD. **(A)** Schematic for model of acute gut-mediated GVHD where donor CD4^+^ T cells respond to host allogeneic peptide presented within donor MHC class II, which in turn drives CD4^+^ T cell expansion in the mesenteric lymph node (mLN). **(B & C)** Representative FACS plots for IFNγ and IL-17A production and Foxp3 expression on TEa cells from the mLN at day 4 post-transfer, with PMA/ionomycin restimulation (B) or directly *ex vivo* (C). Graphs of percentage IFNγ^+^, IL-17A^+^ or Foxp3^+^ TEa cells. Data shown are combined from three replicate experiments (*n* = 10 mice). Statistical analysis was performed between the isotype control and respective cytokine samples using a Mann-Whitney test. *p<0.05, **p≤0.01, ***p≤0.001, ****p≤0.0001.

### Droplet-based, single-cell RNA-seq and computational modelling reveals divergent fates in donor CD4^+^ T cells

To examine donor CD4^+^ T cell differentiation without reliance on pre-selected markers, we employed droplet-based scRNA-seq. TEa cells were recovered from mLN of irradiated BALB/c alloSCT recipients, at days 1, 2, 3 & 4 post-transfer (Figure 2A and Figure S1A & B) – non-transferred control TEa cells were also examined, referred to as day 0 cells. Flow cytometric assessment of sorted TEa cells - pooled from mice at each time-point – revealed as expected rapid, transient upregulation of CD69 and evidence of cell division by CFSE dilution by day 2 (Figure S1B). This was followed on days 3 and 4 by complete loss of CFSE, indicative of dramatic clonal expansion within days 2-3, as well as substantial upregulation of the gut homing integrin, α4β7 in many, but not all TEa cells. These data confirmed that TEa cell activation had occurred in mLN, and suggested emerging heterogeneity. TEa cells were therefore processed for droplet-based scRNA-seq. After excluding poor-quality single-cell transcriptomes (Figure S2A & B), we advanced 22,854 high-quality TEa samples for further analysis. We noted a substantial increase in the average number of genes detected per cell from day 0 through day 1 and 2, which dropped by day 4 (Figure S2A). This was consistent with our previous study in which CD4^+^ T cells more than doubled the number of expressed genes during clonal expansion (9).

**Figure 2.**
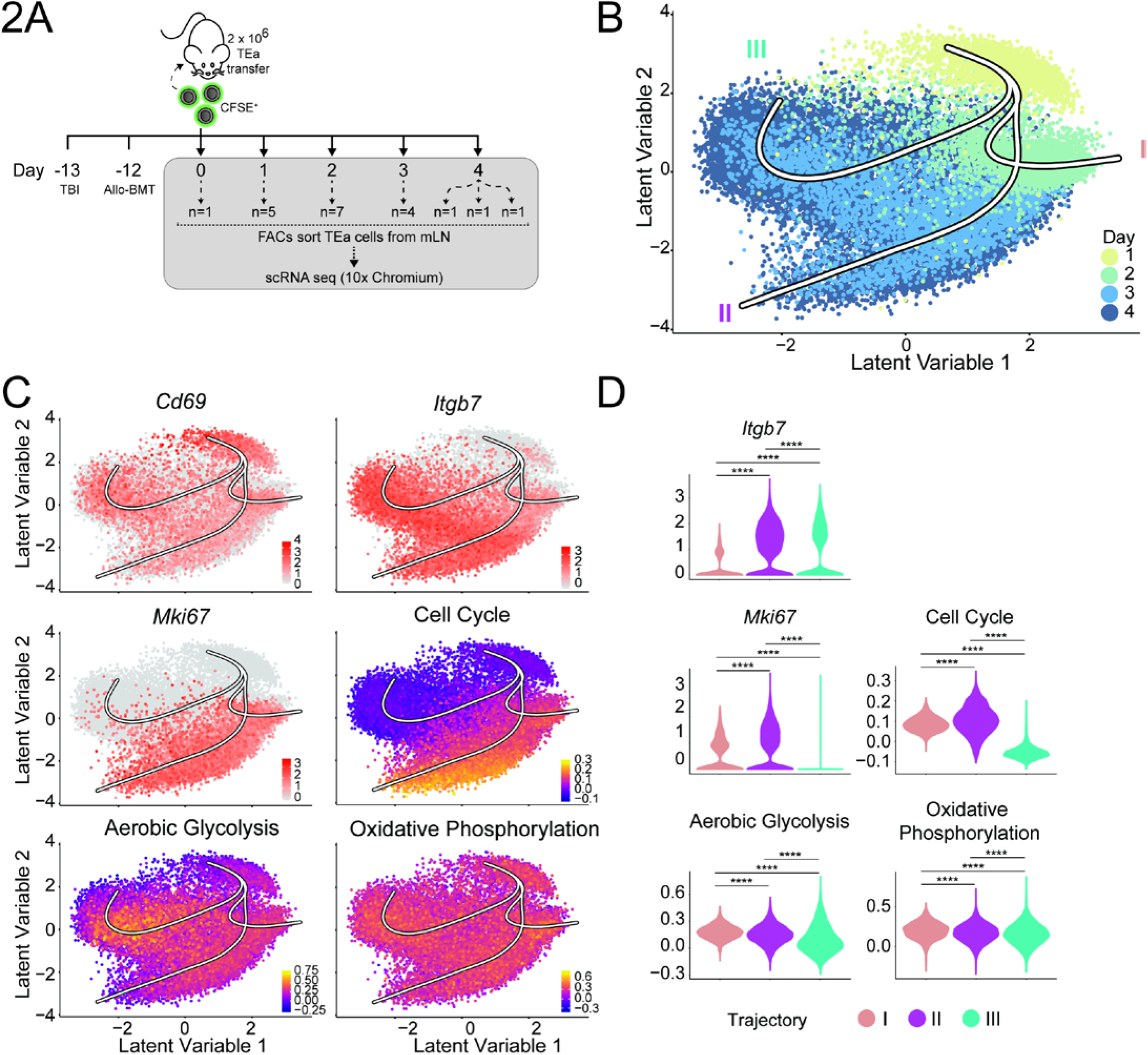
Droplet-based scRNA-seq of TEa cells in the mLN and computational modelling identifies multiple developmental lineages. **(A)** Schematic of scRNA-seq experiment used to study donor CD4^+^ T cell differentiation. TEa cells were recovered from the mLN at various timepoints and cells pooled from multiple mice (n = number of mice pooled; at day 4 samples from 3 mice ran separately) for droplet-based scRNA-seq using the 10x Chromium. **(B)** bGPLVM visualisation of TEa cells from day 1 to day 4 overlaid with the developmental trajectories identified by Slingshot (Trajectory I, II and III). **(C)** Visualisation of *Cd69*, *Mki67* and *Itgb7* expression, or the cell cycle, aerobic glycolysis, and oxidative phosphorylation gene signature scores, in bGPLVM overlaid with trajectories. **(D)** Violin plots showing expression of genes or gene signature scores as described in (C) for each trajectory. Statistical analysis was performed using a Wilcoxon-rank sum test between trajectories. *p<0.05, **p≤0.01, ***p≤0.001, ****p≤0.0001.

Uniform Manifold Approximation and Projection (UMAP) (15) visualisation after Principal Component Analysis (PCA) suggested that day 0 and day 1 TEa cells, whilst relatively homogeneous within their own timepoints, were distinct from each other, and furthermore were distinct from cells assessed at days 2-4 (Figure S2C). In contrast, there was evidence both of transcriptomic overlap between cells from days 2, 3 and 4, as well as of transcriptomic progression from day 2 through to day 4 (Figure S2C). Therefore, we sought to test for potential developmental trajectories within the scRNA-seq data. We employed non-linear probabilistic PCA, termed bGPLVM (9), which embedded the data in low-dimensional space and re-ordered transcriptomes independently of the time-point of capture. Our previous paper had employed bGPLVM on a relatively small number of cells (<1000) (9). It was evident, here, that bGPLVM was scalable to ~25,000 cells. Running bGPLVM iteratively ten times on the dataset yielded similar learned embeddings, indicating the stability of the output (Figure S3). Variability between cells was clearly observable along three latent variables, with interpretable variation also evident in the first two latent variables, thus permitting modelling in two dimensions (Figure 2B). We next sought to infer possible differentiation trajectories based on transcriptomic similarity between cells. We employed Slingshot (16) on the bGPLVM embedding with a start point specified in day 1, and with day 0 omitted due to transcriptomic distance from the rest of the data (Figure S4A & B). Slingshot identified three potential trajectories in the bGPLVM space, two of which (trajectories II & III) progressed through days 2, 3 and 4, while trajectory I appeared to terminate in day 2 (Figure 2B). Consistent with flow cytometric data (Figure S1), *Cd69* expression gradually reduced along trajectories II & III, while *Itgb7*, encoding subunit β7 of integrin α4β7, increased (Figure 2C & D). Most strikingly, trajectory III ceased expression of cell-cycling genes including *Mki67* (encoding Ki-67)(Figure 2C & D) and exhibited lower expression of aerobic glycolysis genes compared to trajectory II. We also employed Slingshot on UMAP of PCs and found four trajectories, two of which largely superimposed upon each other suggesting, as for bGPLVM, three main trajectories (Figure S5A). Two of these progressed through days 2-4, with one largely devoid of cellular proliferation, and the other exhibiting strong expression of *Mki67* and other cycling genes (Figure S5B & C). To further test for trajectories in the data, we employed other computational approaches, including Monocle 2, Velocyto (17), and PAGA (18). In all cases (Figure S6), there appeared one main bifurcation event within days 2-3, leading to two trajectories. Therefore, taken together, our analysis suggested two apparent trajectories had emerged in TEa cells, which differed from each other in expression of genes related to cell cycling and aerobic glycolysis.

### Pro-inflammatory and regulatory effector fates emerge within one trajectory, revealing a second, quiescent *Tcf7*-expressing fate

We next assessed expression of the canonical markers, *Ifng*, *Il17a* and *Foxp3* within our bGLPVM/Slingshot model (Figure 3A), and surprisingly, found them strongly expressed in only trajectory II. In addition, we noted significant expression of *Il17a* and *Il17f* (Figure 3, *Il17f* not shown), in areas shared by trajectory I and II (Figure 3A). These observations pointed to early activation of the *Il17a* promoter at day 2, followed later by upregulation of *Ifng* and *Foxp3*. Most importantly, we saw little evidence of Th1, Th17 and iTreg cell types emerging from different developmental trajectories, either when examined using our previous Th gene modules or when relatively sparse *Ifng*, *Il17a* and *Foxp3* expression levels were imputed using Adaptively-thresholded Low-Rank Approximation (ALRA)(19)(Figure S7). Instead, our unbiased analysis suggested pro-inflammatory and regulatory effector fates had emerged within trajectory II. Importantly, we noted that trajectory III lacked expression of *Ifng*, *Il17a* and *Foxp3* (Figure 3A). Differential gene expression analysis between cells in trajectory III vs II revealed *Tcf7* as a top transcription factor associated with trajectory III, as well as elevated expression of memory-associated *Klf2, Ccr7, Cd27* and *Sell* (encoding CD62L)(Figure 3B & C and Table S1). These observations were also seen in our UMAP/Slingshot model and in a second scRNA-seq experimental repeat (Figure S8). Thus, given an absence of cell cycle activity, lower expression of aerobic glycolysis genes, absence of pro-inflammatory or regulatory gene expression, and expression of memory or stem-like genes including *Tcf7,* we inferred that trajectory III contained quiescent TEa cells that had gone through a clonal burst in mLN, but had not acquired any effector function. To test predictions from our transcriptomic models, we assessed mLN TEa cells at day 4 post-transfer by flow cytometry. As expected, all TEa cells had upregulated the Th1-associated lineage transcription factor, T-bet (Figure 4A). We also observed a clear bifurcation amongst these cells in expression of the *Tcf7*-encoded protein, TCF1 (Figure 4A), with one population of cells expressing higher levels of TCF1 and lower levels of T-bet than its counterpart (Figure 4A). We also noted that direct *ex vivo* expression of IFNγ or IL-17A was substantially reduced in TCF1^hi^ cells relative to TCF1^lo^ counterparts within the same mouse. Strikingly, Foxp3 expression was absent in TCF1^hi^ cells relative to TCF1^lo^ cells (Figure 4A), and *in vitro* re-stimulation did not recover expression of these molecules, (Figure 4B). Together these data validated our prediction from scRNA-seq modelling of the differentiation of a major population of allo-reactive CD4^+^ T cells in mLN, which were quiescent and marked by high expression of TCF1.

**Figure 3.**
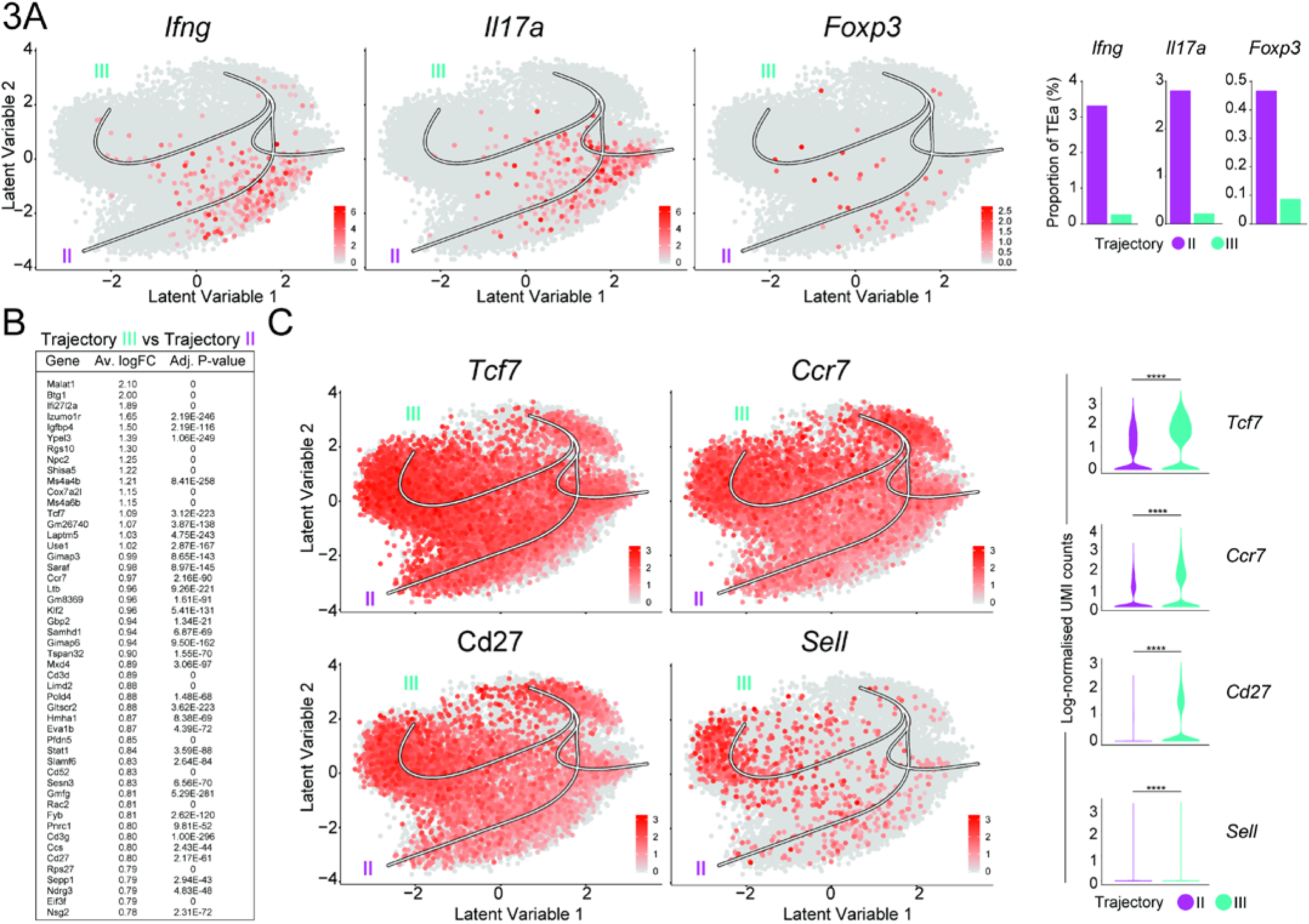
*Ifng*, *Il17a* and *Foxp3* expressing fates emerge along one trajectory, identifying a second *Tcf7*-expressing fate. **(A)** bGPLVM visualisation of TEa cells expressing *Ifng*, *Il17a* and *Foxp3* with Slingshot trajectories overlaid. Bar graphs show the proportion of cells within Trajectory II or III that express *Ifng*, *Il17a* or *Foxp3*. **(B)** Top 50 genes upregulated in Cluster 0 in Trajectory III compared to Cluster 3 in Trajectory II, identified using the “FindMarkers” function in Seurat. Average log fold change (Av.logFC) and Bonferroni adjusted P-value (Adj. P-value) are shown. **(C)** bGPLVM visualisation of TEa cells expressing *Tcf7*, *Ccr7*, *Cd27* and *Sell* with Slingshot trajectories overlaid. Violin plots show the level of expression for each gene in Trajectory II and III. Statistical analysis was performed using a Wilcoxon-rank sum test between trajectories. *p<0.05, **p≤0.01, ***p≤0.001, ****p≤0.0001.

**Figure 4.**
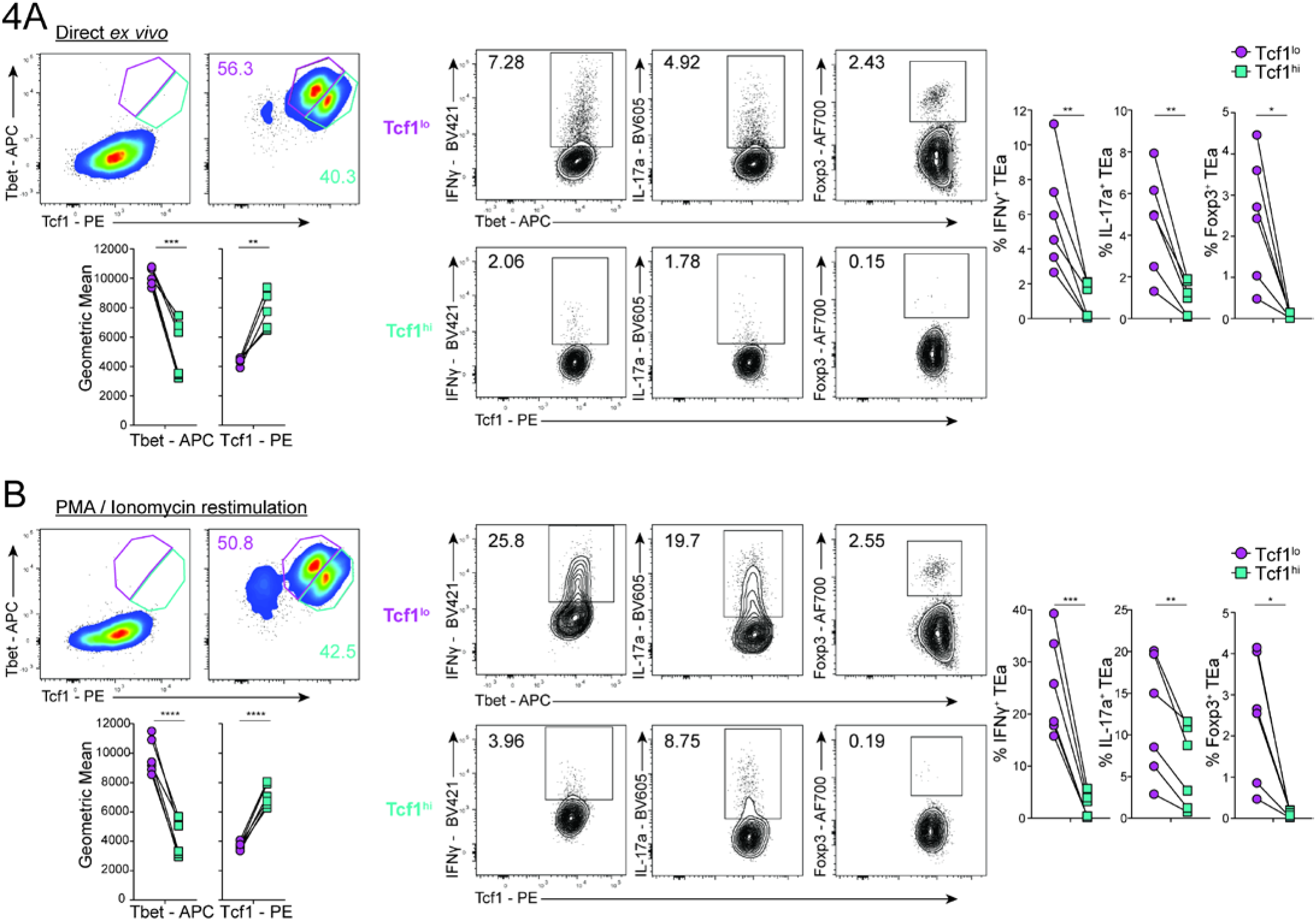
TEa cells exhibit heterogeneity in the mLN. **(A & B)** Representative FACS plots showing the expression of Tbet and Tcf1 in TEa cells at day 4 post-transfer in the mLN directly *ex vivo* (A) or after restimulation with PMA/ionomycin (B). Graphs show the geometric mean fluorescence intensity of the Tcf1^lo^ (purple) or Tcf1^hi^ (turquoise) population. The expression of IFNγ, IL-17A and Foxp3 for the Tcf1^lo^ (purple) or Tcf1^hi^ (turquoise) population is also shown. Graphs show the percentage of IFNγ^+^, Il-17A^+^ or Foxp3^+^ TEa cells for the Tcf1^lo^ or Tcf1^hi^ population. Data shown are combined from two independent experiments (*n* = 3 mice / experiment). Statistical analysis was performed using a paired *t*-test. *p<0.05, **p≤0.01, ***p≤0.001, ****p≤0.0001.

### *Tcf7*^hi^ allo-reactive CD4^+^ T cells change minimally during migration from mLN to gut

Although TCF1^hi^ TEa cells within the mLN failed to express Foxp3, or pro-inflammatory cytokines, IFNγ and IL-17A, scRNA-seq predicted their capacity to migrate to the gut due to expression of the integrin gene *Itgb7* (Figure 2C). To test this, and to explore the developmental relationships between mLN and gut-migrating TEa cells, we conducted a third scRNA-seq experiment, examining TEa cells at day 5, both in mLN and in the intra-epithelial lymphocyte (IEL) fraction of the gut (Figure 5A). TEa cells were readily recovered from lamina propria (LP) and IEL fractions from the gut. However, given the longer protocol required to isolate cells from LP versus IEL, and the potential to interfere with transcriptome fidelity, we confined scRNA-seq analysis to IEL TEa cells. To relate day 5 data with our previous mLN dataset, we repeated assessments of days 0 and 4 mLN. Finally, to control for possible technical variation in IEL induced by the isolation protocol, we treated day 5 mLN cells with and without the IEL-isolation pre-digestion buffer. After scRNA-seq, and quality control as before (Figure S9A), we assessed 21,632 cells across all samples. Firstly, we noted no effect of the IEL isolation buffer on transcriptomes (Figure S9B), indicating that direct comparison of cells from mLN and IEL was possible. Next, unsupervised clustering of day 5 IEL cells, revealed four main clusters (0, 1, 2 & 3, and minor cluster 4), with *Ifng, Il17a* and *Foxp3* upregulated across a broad area in Clusters 0, 1 & 3 but not 2 (Figure S9C). The frequency of TEa cells expressing pro-inflammatory cytokine genes was substantially elevated compared to our earlier experiments in mLN, with ~15% and ~40% of TEa cells in Clusters 0, 1 & 3 expressing *Il17a* or *Ifng* respectively (Figure S9C). Pronounced expression also revealed that patterns for *Ifng* and *Il17a* expression were not identical in IEL TEa cells. While *Ifng* was relatively uniform in expression across Clusters 0, 1, & 3, *IL17A* was focussed in specific areas. However, a clear distinction between Th1 and Th17 could not be drawn transcriptomically, consistent with previous data that allo-reactive CD4^+^ T cells can co-express IFNγ and IL-17A at protein level. Interestingly, *Foxp3* was not tightly confined to a particular area in IEL cells, suggesting that iTregs were transcriptomically varied, and could not be partitioned into a specific lineage separate from those expressing pro-inflammatory cytokines. Together, these data reveal that even after priming in mLN and migration to the gut, allo-reactive effector CD4^+^ T cells remain transcriptomically similar to each other, regardless of their pro-inflammatory or regulatory phenotype. Unsupervised clustering of IEL TEa cells also revealed Cluster 2, which completely lacked expression of *Ifng, Il17a* and *Foxp3*, but was elevated for *Tcf7*, *Ccr7* and *Cd27* expression relative to other clusters (Figure S9C). Cluster 4 contained a small population of cells that could not be reliably analysed (Figure S9D). Thus, our assessment of TEa cells in the gut revealed the presence of both effector and quiescent cell states, mirroring those seen in mLN.

**Figure 5.**
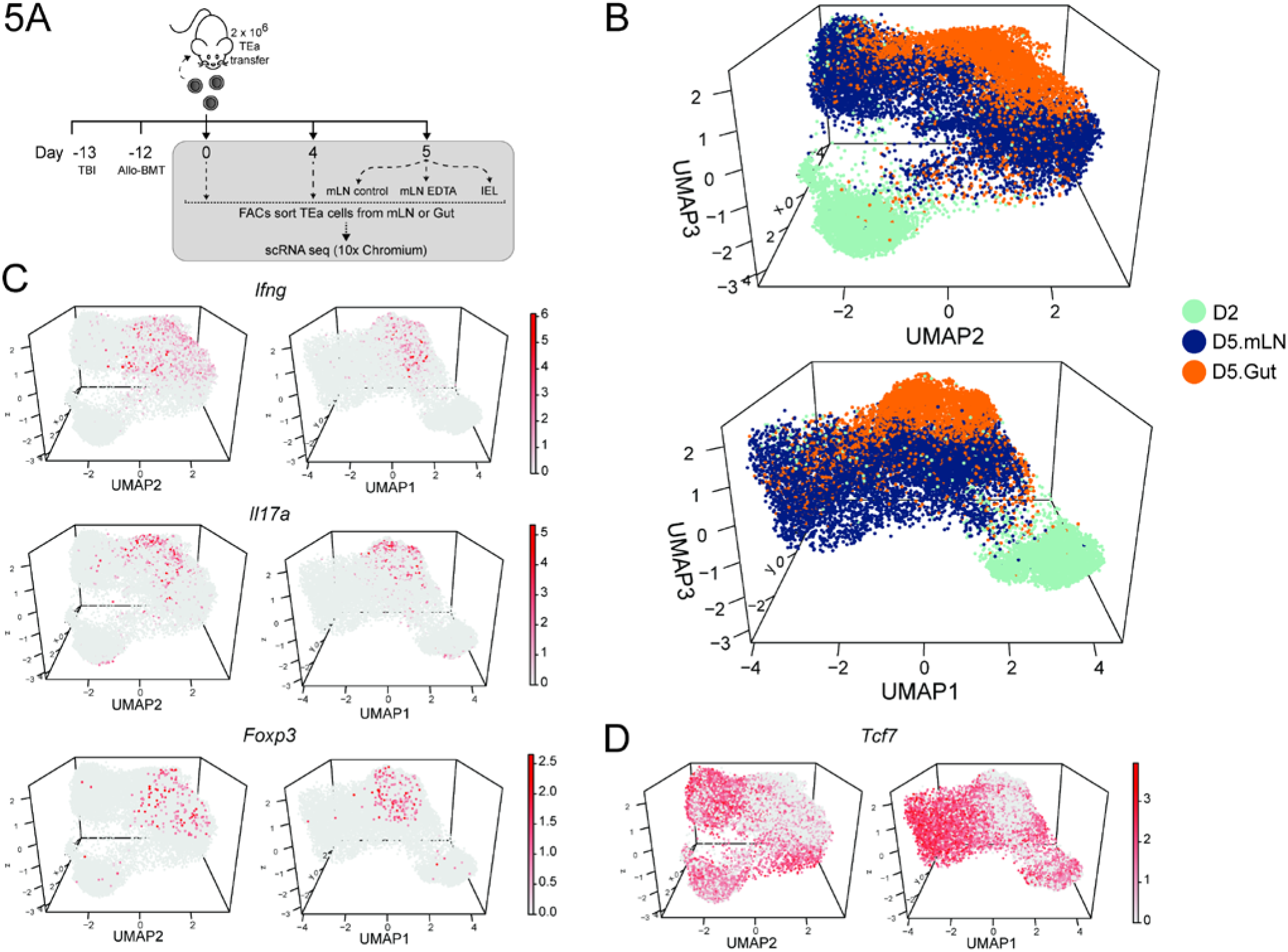
*Tcf7*^hi^ TEa cells migrate from the mLN to the gut and remain transcriptomically similar. **(A)** Schematic of scRNA-seq experiment used to study the developmental relationships between TEa cells from the mLN and the gut. **(B)** Two perspectives of 3-Dimensional UMAP visualisation produced from 30 scVI latent variables representing all samples from day 2 - day 5. Only day 2, day 5 gut, and day 5 mLN samples are shown for clarity. **(C)** *Ifng*, *Il17a*, and *Foxp3* expression in the same representation as (B). **(D)** *Tcf7* expression in the same representation as (B) and (C).

To determine molecular relationships between TEa cells in mLN and gut, we integrated our datasets across all time-points and organs using single-cell Variational Inference (scVI) (20), which accounted for possible batch effects across independent experiments, and provided a temporal atlas of differentiation for allo-reactive CD4^+^ T cells in the gut (Figure S10A-B). Output from scVI was first visualised via 3-dimensional UMAP. Day 0 and day 4 mLN TEa cells, regardless of experiment, occupied the same space as their time-point counterparts, suggesting that any technical variation from different experiments, sequencing platforms, and protocols had been removed. Next, we noted that day 0 and day 1 cells existed in discrete transcriptional states, separate from each other and from day 2-5 cells (Figure S10A-B). This suggested TEa cells had undergone rapid and uniform change upon initial exposure to allo-antigen in mLN, with potential intermediate cellular states not captured by scRNA-seq assessment at a single time-point 24 hours after transfer. Similarly, we noted few transcriptomic intermediates between day 1 and day 2, suggesting again further uniform, rapid change during the second 24-hour period. Differential gene expression analysis between consecutive days revealed gene families associated firstly with ribosomal processes, then cellular division were upregulated during the first 48-hour period of allo-antigen exposure (Table S2), consistent with initiation of clonal expansion. Next, we noted from day 2-5 in mLN, a substantial, gradual increase in heterogeneity, with modest effector molecule expression, such as *IFNg, IL-17a* and *Foxp3* expression confined to one space, with quiescent *Tcf7^hi^* cells occupying a separate space (Figure 5C-D). Importantly, from our integrated model we inferred further transcriptomic change as effector cells migrated from mLN to gut, including for example, upregulation of *Csf2* (encoding GM-CSF), Tr1-associated immune-suppressive *Il10,* and increased expression of *IFNg, IL-17A* and *Foxp3* (Figure S10C). This suggested differentiation into *IFNg^+^ or IL-17A*^+^ pro-inflammatory effectors, Foxp3^+^ iTreg cells or IL-10^+^ Tr1 cells had not been completed in secondary lymphoid tissue and instead occurred progressively during and after migration to the gut. In contrast, we noted substantial transcriptomic overlap between *Tcf7^hi^* cells in mLN versus IEL. Moreover, once *Tcf7*^hi^ cells had emerged in mLN by day 3, their phenotype altered very little either over the following 2 days or during migration to the gut (Figure 5D). These data suggested that *Tcf7^hi^* cells remained transcriptomically stable across different tissues, while effector cells, including those expressing *Ifng, Il17a,* or *Foxp3*, underwent progressive transcriptomic change over this period.

Finally, using our scVI model, we sought to examine how quiescent, allo-reactive CD4^+^ T cells developed in mLN during acute gut GVHD. Trajectory inference, for example using Slingshot on bGPLVM or UMAP embedding, assumed developmental changes in cells are gradual enough to capture intermediate states. However, while emergence of effector cells appeared gradual in the mLN and during migration to the gut, this was less clear for emergence of *Tcf7*^hi^ cells. Transcriptomic intermediate states were apparent between *Tcf7*^hi^ and *Tcf7*^lo^ cells by days 4-5 (Figure 5D), consistent with possible linear transitions from effector to memory. However, at day 2 prior to effector differentiation and clonal expansion, small numbers of cells appeared in the quiescent, low cell-cycling *Tcf7*^hi^ population (Figure 5D), suggesting an alternative mechanism of development unlinked to effector differentiation. Thus, scRNA-seq analysis revealed that some quiescent, *Tcf7*^hi^ allo-reactive CD4^+^ T cells had emerged rapidly within 48 hours of allo-presentation in mLN, with developmental intermediates being difficult to capture in our experimental design. In summary, our integrated atlas of allo-reactive CD4^+^ T cell differentiation (Movie S1), revealed the emergence of pro-inflammatory, regulatory, and quiescent cell states within secondary lymphoid tissue, which emerged rapidly and evolved to differing degrees during migration to the gut.

### TCF1^hi^ quiescent allo-reactive CD4^+^ T cells retain a capacity to mount effector responses *in vivo*

We sought to examine the functional potential of quiescent TCF1^hi^ TEa cells during acute gut GVHD. In particular, we sought to test the hypothesis that TCF1^hi^ TEa cells could eventually give rise to an effector response within the gut, and re-generate themselves. Firstly, we noted that although these cells expressed high levels of *Pdcd1* (encoding PD-1), they did not express other co-inhibitory markers often associated with T cell exhaustion, suggesting they might still be responsive to re-activation (Figure S11A). Secondly, as for naïve cells, *Tcf7*^hi^ cells expressed high levels of both *Il6st and Il6ra,* which are required for classical IL-6 signalling that promotes CD4^+^ T cell responses in acute gut GVHD (Figure S11B). These data suggested that *Tcf7*^hi^ cells retained the capacity to respond to allo-antigen. Therefore, we designed a cell-sorting strategy for *Tcf7*^hi^ cells based on our bGPLVM/Slingshot scRNA-seq model, in which trajectory III cells expressed much lower levels of *IL2ra* (encoding CD25) and the Th1-associated chemokine receptor gene, *Cxcr6* compared with trajectory II effector cells (Figure S12A). Consistent with this, flow cytometric assessment of day 4 mLN revealed that IFNγ/IL-17A production and Foxp3 expression was confined to CD25^+^ TCF1^lo^ TEa cells (Figure S12B). Hence, we sorted clonally-expanded (CFSE^lo^) CD25^−^ CXCR6^−^ TEa cells from mLN at day 4 post-transfer, as well as CD25^+^ and/or CXCR6^+^ counterparts, and transferred these or naïve TEa control cells into irradiated BALB/c alloSCT recipients (Figure 6A & Figure S12C). When assessed 4 days later, recipients of TCF1^+^ TEa cells harboured as many cells as those receiving naïve TEa cells (Figure 6B). In contrast, recipients of TCF1^−^ TEa cells, albeit transferred with 50% lower numbers, harboured fewer Tea cells (~10% of original input, compared to 40-60% of input for naïve and TCF1^hi^ cells). While naïve TEa cells diverged to give rise to both TCF1^hi^ and TCF1^lo^ cells, TCF1^+^ TEa cells were less able to re-generate the TCF1^hi^ phenotype in a secondary transplant, and TCF1^−^ TEa cells were almost incapable of doing so (Figure 6C). TCF1^+^-derived TEa cells were capable of expressing IFNγ, IL-17A or Foxp3 directly *ex vivo* and after re-stimulation (Figure 6D & E), with IL-17A expression increased relative to primary responses by naïve cells. Together, these data indicated that TCF1^+^ allo-reactive CD4^+^ T cells, though quiescent during a primary response in mLN, retained a capacity to mount effector responses, but were poor at regenerating the TCF1^hi^ phenotype a second time. Thus, we reveal during acute gut GVHD a quiescent, gut migratory, allo-reactive CD4^+^ T cell state that retains effector potential.

**Figure 6.**
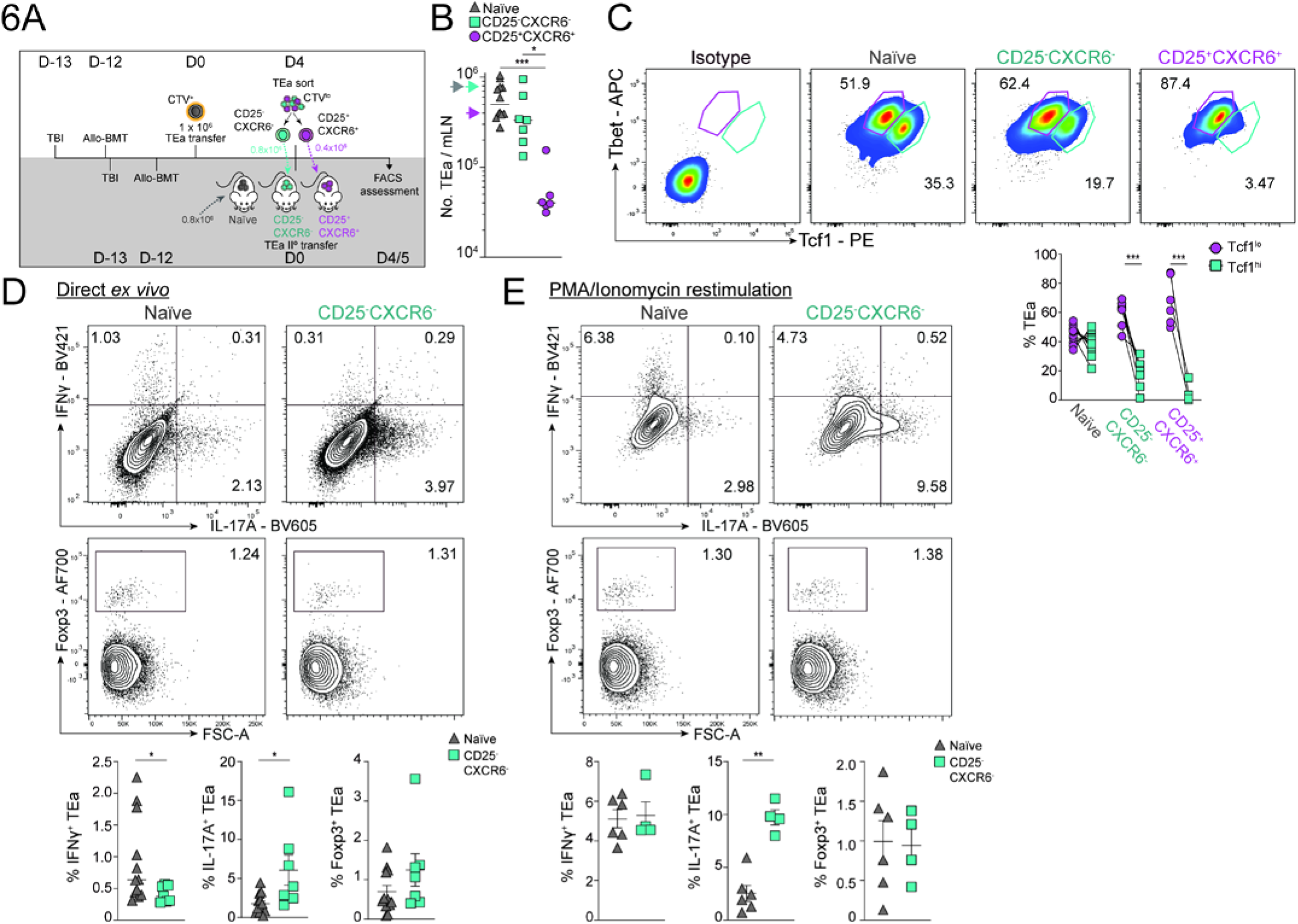
TCF1^hi^ quiescent allo-reactive CD4^+^ T cells retain a capacity to mount effector responses *in vivo*. **(A)** Schematic of secondary TEa transfer experiment. On day 4 of primary transplant, CD25^−^CXCR6^−^ or CD25^+^CXCR6^+^ TEa were FACS isolated from mLN. 0.8 × 10^6^ CD25^−^CXCR6^−^ and 0.4 × 10^6^ CD25^+^CXCR6^+^ TEa (or control 0.8 × 10^6^ Naïve TEa) were transferred into recipient BALB/c mice that had received total body irradiation 13 days prior and a bone marrow transplant 12 days prior. FACS assessment of mLN TEa cells was performed on day 4 or day 5 post-secondary transfer. **(B)** Absolute numbers of TEa cells in the mLN at day 4/5 post-secondary transfer. Arrows along y-axis denote the number of TEa cells that were transferred per mouse for each group on day 0 (0.8 × 10^6^ Naïve TEa (*grey*) and CD25^−^CXCR6^−^ TEa (*turquoise*), 0.4 × 10^6^ CD25^+^CXCR6^+^ TEa for (*purple*)). **(C)** Representative flow cytometry plots showing the expression of Tbet and Tcf1 on TEa cells from the mLN at day 4/5 post-secondary transfer. Graph shows % of TEa cells that are Tcf1^lo^ or Tcf1^hi^ in each group. **(D & E)** Representative FACS plots showing IFNγ, IL-17A and Foxp3 expression on TEa cells after secondary transfer on day 4/5 from the mLN directly *ex vivo* (D) or post-restimulation with PMA and ionomycin (E). Graphs show the percentage of IFNγ^+^, IL-17A^+^ and Foxp3^+^ TEa for each group. Data shown are combined from two independent experiments showing similar results (B-D; Naïve *n =* 12, CD25^−^CXCR6^−^ *n=* 7, CD25^+^CXCR6^+^ *n* = 6) or from one experiment only (E; Naïve *n =* 6, CD25^−^CXCR6^−^ *n=* 4). Data are represented as median (B) or mean ± SEM (C - E). Statistical analysis was performed using one-way ANOVA with multiple comparisons (B), Paired *t* test (C), or a Mann-Whitney test (D, E). *p<0.05, **p≤0.01, ***p≤0.001, ****p≤0.0001.

## Discussion

Although alloSCT is an established curative therapy for a range of haematological malignancies, a major limitation is acute GVHD, where alloreactive naive donor T cells differentiate into pro-inflammatory effectors that damage the GI tract, liver and skin (1). Cytokines such as IL-17A, IFNγ and GM-CSF produced by allo-reactive Th1 and Th17 cells in the GI tract promote disease (21), while IL-10 produced by Tr1 and iTreg cells can protect (22). An important goal in alloSCT is to preserve GVL effects while reducing GVHD. Key to this endeavour is a consideration of the spatial and temporal differences between GVL and GVHD. While GVL exerts beneficial effects in primary and secondary lymphoid organs, acute and lethal GVHD often occurs in the GI tract. By understanding CD4^+^ T cell differentiation after alloSCT, we may define new strategies to block pathogenic cellular states and encourage protective ones. Although CD4^+^ T cell differentiation has been explored at genome-scale in infection, auto-immune and allergy models (7–9), extrapolating to alloSCT remains challenging. For example, in the alloSCT setting, alloantigen is ubiquitous and constant, while pathogen-derived antigen may be more dynamic or transient. Secondly, alloSCT often features profound lymphopenia unlike other models. Hence, we specifically examined transcriptome dynamics of T helper cell differentiation in the GI tract of mice after alloSCT.

By sampling transcriptomes from thousands of alloreactive CD4^+^ T cells of a single specificity across lymphoid and nonlymphoid gut-associated tissue, we successfully detected cellular states expressing canonical Th1/Th17 cytokine genes, *Ifng* and *Il17a*, and the regulatory genes, *Foxp3* and *Il10* – at frequencies similar to that observed by flow cytometry. Importantly, however, unbiased clustering and trajectory inference tools suggested substantial similarity between the transcriptomes of these effector subsets, particularly in lymphoid tissue but also in the gut. Subtle differences became more evident amongst gut trafficked TEa cells that had stopped proliferating, with *Ifng* expressed more uniformly than either *Il17a* or *Foxp3*. Given that TEa cells can upregulate T-bet and IFNγ but not IL-17A or Foxp3 in the complete absence of MHCII-presentation (3, 4), our data are broadly consistent with a Th1-like state being a default program in the gut, that may be countered by alloantigen presentation via MHCII towards iTreg or Th17-like states. Nevertheless, our main inference from transcriptome dynamics was that pro-inflammatory states were not readily distinguished from immune-suppressive iTreg states. We propose a model for development of Th1, Th17 or iTreg-like states during alloSCT controlled by specific micro-anatomical, T cell extrinsic factors, such as access to MHCII-presentation on different types of APC, or exposure to IL-6 and IL-12-signalling. Given the apparent continued maturation of pro-inflammatory TEa cells as they migrated from mLN to the gut, this raises the question of whether peripheral tissue signals in the gut, such as local cytokine-signalling or unique cell-cell interactions contribute to this process. We speculate that spatial transcriptomic assessment of gut-located TEa cells will shed light on this matter. One further implication of our data is that conversion of emerging pro-inflammatory CD4^+^ T cells into protective, non-pathogenic iTreg cells in the gut may be feasible, since developmental pathways are similar between the two. Conversely, our data also support iTreg-based therapies with appropriate mitigation for the effects of reversion to pro-inflammatory states *in vivo* (23).

We and others have previously studied Th17 biology via fate-mapping using *IL-17a* promoter-driven, Cre-mediated fluorescent tagging of cells (6, 24, 25). This binary approach is powerful, but does not differentiate between cells that might transiently express *Il17a* compared with those exhibiting prolonged expression. In our scVI transcriptomic model, we noted early transient expression of *Il17a* and *Il17f* at day 2, which disappeared only to re-appear in some cells at day 5 in the gut. This raises the possibility that CD4^+^ T cells can transiently express *IL-17a* in mLN without becoming *bona fide* Th17 cells. Hence, we propose that studies employing permanent marking of cells using Cre-LoxP systems should be interpreted with possible transient expression taken into account.

Our unbiased, single-cell genomic approach identified an unexpected, apparent trajectory characterised by TCF1^hi^ expression, rapid shutdown of cellular proliferation, a lack of pro-inflammatory or immune-regulatory gene expression, an ability to migrate to the gut, and a capacity to mount a secondary recall response despite modest expression of select checkpoint blockade molecules. Based on these observations we infer TCF1^hi^ TEa cells to be generally quiescent, memory or stem-like cells that emerged very rapidly during alloSCT. Transcriptomic modelling suggested that *Tcf7*^hi^ cells could under certain circumstances arise from the cytokine-expressing effector lineage at day 3 or 4, which would be consistent with a linear model in which effector cells give rise directly to memory-like cells (26, 27). However, we also noted rare instances of *Tcf7*^hi^ cells emerging at day 2 post-transfer, as clonal expansion was beginning and effector differentiation had yet to occur. We did not detect transcriptomic intermediates between this distal state and more naïve cells, either because such states do not exist, or because our study was not designed to detect such rare, transient events. Given that trajectory inference from scRNA-seq data tends to rely upon, indeed assume, gradual transcriptomic change, it is likely that very rapid state changes cannot be mapped using this approach. Our data does not resolve the extent to which TCF1^hi^ CD4^+^ T cells emerge via gradual linear effector-memory transition versus a more rapid, branching process via asymmetric cell division, which was recently reported for similar cells during a respiratory virus infection model (28), and for CD8^+^ T-cells during LCMV infection (12). We speculate, given the presence of rare “pioneer”-like *Tcf7*^hi^ cells as well as apparent transcriptomic intermediates, that both these developmental pathways may operate during alloSCT. New research tools are required to quantify the relative use of these mechanisms *in vivo*.

One question to consider from our transcriptomic modelling is how separate states, loosely referred to as “effector” or “quiescent”, emerge amongst clonal T cells during alloSCT. Heterogeneity amongst clonal TEa cells could have been induced via various non-mutually exclusive mechanisms, including asymmetric cell division (28), differential APC engagement and differential access to early local cytokine signals. Given the stark difference in CD25 expression between trajectories, it appears feasible that IL-2-signalling promotes effector function at the expense of quiescence. A recent study revealed a reciprocal relationship in production and receipt of IL-2 controlled fate bifurcation in CD4^+^ T cells (29). Hence, we speculate that a similar mechanism might be acting during alloSCT. In our model, we previously reported that colonic CD103^+^ dendritic cells played a crucial role in amplifying acute GVHD (4). It is possible that naïve CD4^+^ T cells that failed to access these APC may have been programmed towards the quiescent, TCF1^hi^ state. As part of this scenario, given that our model is characterised by profound lymphopenia, it is possible that homeostatic proliferation, perhaps via IL-7 signalling, might have partly contributed to the proliferation and stabilisation of a quiescent cellular state. However, given that donor allo-antigen presentation via MHCII is important for supporting clonal expansion in this model, exposure to diverse donor APC may contribute to heterogeneity in CD4^+^ T cell differentiation.

Our experiments revealed that clonally expanded TCF1^hi^ CD4^+^ T cells, which had exhibited no effector function and had shut down cellular proliferation, were capable of mounting a pro-inflammatory or regulatory response at a later date, as shown during secondary transplantation. This suggests that after alloSCT, the gut could be populated with quiescent alloreactive T cells that could influence disease outcome at a later timepoint. Despite their capacity to mount either pro-inflammatory or regulatory responses, we view these cells as an opportunity to lodge immune-suppressive, possibly tissue-resident CD4^+^ T cells within the gut that could contribute to disease prevention after alloSCT. However, the potential for these to produce pathogenic cytokines would clearly need to be addressed. In summary, our examination of transcriptome dynamics during alloSCT not only highlighted developmental relationships between effector CD4^+^ T cells, but also uncovered the existence of a quiescent, memory-like state that exhibits functional potential *in vivo*. It will be important next to interrogate our experimentally-derived observations in patient samples after alloSCT.

## Methods

### Mice

C57BL/6J and BALB/c mice were purchased from the Animal Resources Centre (Canning Vale, Australia), transgenic TEa (Vα2^+^, Vβ6^+^, CD45.1^+^, CD90.1^+^) mice were bred in-house (4, 5). All mice were female and aged between 6 – 12 weeks of age and were maintained under specific pathogen-free conditions within the animal facility at QIMR Berghofer Medical Research Institute.

### Bone marrow transplantation

Balb/c mice were transplanted as previously described (4). Briefly, BALB/c mice received 900 cGy total body irradiation (TBI; ^137^Cs source at 84 cGy/min) on day −13. On day −12, Balb/c mice were transplanted with 10 × 10^6^ bone marrow cells and 2 × 10^5^ FACs purified T cells from C57BL/6J donor mice. On day 0, recipient BAL B/c mice were injected intravenously with 1-2 × 10^6^ FACS sorted TEa T cells (Vβ6^+^Vα2^+^). Cell Trace™ CFSE Cell Proliferation (Life Technologies) and Violet Proliferation Dye 450 (VPD450; BD Biosciences) staining were performed according to manufacturer’s protocol. For secondary transfer of TEa cells, CD25^−^ CXCR6^−^ TEa cells or CD25^+^CXCR6^+^ TEa cells were FACS isolated on D4 from the mLN and then injected intravenously into BALB/c mice that had received total body irradiation 13 days prior and transplanted with bone marrow cells and purified T cells from C57BL/6J mice 12 days prior. For reasons of cell availability post-sort, 0.8 × 10^6^ CD25^−^CXCR6^−^ or 0.4 x10^6^ CD25^+^CXCR6^+^ TEa cells were transferred into respective groups of mice, and 0.8 × 10^6^ naïve TEa cells transferred as a reference control.

### Cell isolation

#### Spleen

Spleens were collected and homogenised through a 100 μm cell strainer to generate a single cell suspension. Red blood cells were then lysed using RBC Lysing Buffer Hybri-Max™ (Sigma-Aldrich). Splenocytes were then washed with RPMI media (Gibco) prior to proceeding with staining for flow cytometry assessment.

#### Small intestine and colon

Intraepithelial lymphocytes were isolated from the small intestine and colon of mice using the Lamina Propria Dissociation Kit, Mouse (Miltenyi Biotec), according to the manufacturer’s protocol.

#### Flow cytometry

Cells were assessed for viability by staining with 7AAD (Sigma-Aldrich) or using a LIVE/DEAD™ Fixable Aqua Dead Cell Stain Kit (Life Technologies), according to manufacturers’ protocol. Prior to staining, cells were incubated with antibodies against CD16 and CD32 (2.42G) to block Fc receptors. For surface staining, cells were incubated with the following antibodies for 20 minutes at 4°C: CD4 (RM4-5) – PerCPCy5.5, CD4 (RM4-5) – PE Dazzle 594, Vα2 (B20.1) – APC Cy7, CD45.1 (A20) – PeCy7, CD69 (H1.2F3) – PB, α4β7 (KATK32) – PE, CD25 (PC61) – PECy7, CXCR6 (SA051D1) – APC, Armenian hamster IgG isotype control PB, rat IgG2a isotype control – PE, rat IgG1k isotype control – PECy7, rat IgG2bk isotype control – APC (all Biolegend). For assessment of intracellular cytokine production and transcription factor expression, cells were incubated with or without ionomycin (500 ng/ml) and PMA (50 ng/ml) for 4 hours at 37°C, brefeldin-A (5 μg/ml) was added to cells after 1 hour of incubation. Intracellular staining was then performed using the eBioscience™ Foxp3/Transcription Factor Staining Buffer Set with the following antibodies: IFNγ (XMG1.2) BV421, IL-17A (TC11-18H10.1) – BV605, rat IgG1k isotype control – BV421, rat IgG1k isotype control –BV605, rat IgG2a isotype – AF700, mouse IgG1k isotype control – APC from Biolegend; Tcf1 (C63D9) – PE, rabbit IgG isotype control – PE from Cell Signaling Technology; Tbet (4B10) – APC, Foxp3 (FJK-16S) – AF700 from eBioscience. Samples were acquired on a LSRII Fortessa Analyser (BD Biosciences) and data analysed using FlowJo software (Treestar).

#### Single-cell RNA capture and sequencing

Three independent experiments were performed for sequencing: termed mGVHD1, mGVHD2, and mGVHD3. TEa cells were isolated by flow cytometry into a 1% BSA/PBS buffer. Approximately 8,000 cells were loaded per channel onto a Chromium controller (10x Genomics) for generation of gel-bead-in-emulsions. Sequencing libraries were prepared using Single Cell 3’ Reagent Kits v2 (mGVHD1 and mGVHD2) or v3.0 (mGVHD3) (10x Genomics) and either sequenced on an Illumina NextSeq550 (mGVHD1 and mGVHD2) or converted using the MGIEasy Universal Library Conversion Kit (BGI) before sequencing on a MGISEQ-2000 instrument (BGI; mGVHD3).

#### Processing of scRNA-sequencing data

FASTQ files were processed using “cellrange count” pipeline from Cell Ranger version 2.1.0 and 3.0.2 (10x Genomics) with 10x mouse genome 1.2.0 release as a reference. For BGI FASTQ files (mGVHD3), it was made compatible with Cell Ranger by reformatting file names and FASTQ headers using code from https://github.com/IMB-Computational-Genomics-Lab/BGIvsIllumina_scRNASeq (30).

#### Quality control of scRNA-seq data

Cells outside the thresholds of 200 – 6,000 expressed genes and up to 15% mitochondrial content were removed. Further filtering was performed after unsupervised clustering of cells, where clusters of cells with low Cd3 and Cd4 expression were removed. Cells that expressed Cd8a were globally removed from downstream analysis. Only genes expressed in 3 or more cells were considered.

#### Data transformation

scRNA-seq data was normalised using “NormalizeData” function from Seurat v2.3.4 (31) where UMI counts for each gene from each cell were divided by the total UMI counts from that cell, multiplied by the scale factor of 10,000 and natural log-transformed. Total UMI content and mitochondrial content per cell were considered unwanted sources of variation and were removed by individual linear regression. Final residuals were then scaled to have mean feature expression of 0 and variation of 1 across cells.

#### Feature selection and dimensionality reduction

Highly variable genes (HVGs) were identified using “FindVariableGene” function from Seurat with default parameters. For each subset of data used for dimensionality reduction, HVGs were computed individually and used as an input, unless otherwise specified. Number of HVGs used in each analysis is noted in the figure legends.

Principal component analysis (PCA) dimensionality reduction was performed using “RunPCA” function from Seurat and the computed PCs were used to generate uniform manifold approximation and projection (UMAP) of scRNA-seq data using “RunUMAP” function from Seurat. Number of PCs used in each analysis is noted in the figure legends.

Bayesian Gaussian Processes Latent Variable Modelling (bGPLVM) dimensionality reduction was performed using GPfates v1.0.0 (9) where datasets containing all genes after initial gene filtering step were used as input. Up to five latent variables were considered.

Integrated dimensionality reduction of mGVHD2 and mGVHD3 datasets was performed using scVI v0.4.1 (20). Each experiment was identified as separate batch. All parameters were kept at default except: up to 30 latent variables were considered, 2 hidden layers were used for encoder and decoder neural networks and up to 100 epochs were used to train the model. The computed latent variables were used as an input to generate UMAP using “RunUMAP” from Seurat, or using the umap-learn v0.3.5 python package (15).

#### Unsupervised clustering

“FindClusters” function from Seurat was used to perform unsupervised clustering of cells. The resolution parameter and the number of PCs or variables used in each analysis are noted in the figure legends.

### Trajectory inference

#### Slingshot

Trajectories were inferred through the mGVHD2 dataset using Slingshot v0.99.12 (16). Slingshot requires clusters of cells and embeddings for those cells as input. We used bGPLVM latent variables 1 and 2, and also performed unbiased clustering based on these variables. A semi-supervised approach was taken whereby a cluster with high proportion of day 1 cells were specified as a starting point (Figure S4A).

#### Monocle

Trajectories were inferred through the mGVHD2 dataset using Monocle v2.8.0 (32). PCA dimensionality reduction was performed and the first 10 PCs were used as an input for unsupervised clustering using “plot_pc_variance_explained” and “clusterCells” functions, where number of clusters was specified (n=6). Differential gene expression analysis was performed between clusters using “differentialGeneTest” function and the list of significant genes (qval < 0.01) was used as an input to order the cells using “orderCells” function.

#### PAGA

Trajectories were inferred through mGVHD2 dataset using SCANPY v1.4.4, which includes the PAGA trajectory inference algorithm (18). PCA dimensionality reduction was performed using the “scanpy.tl.pca” function with ARPACK SVD solver to aid computation. A neighbourhood graph was computed via “scanpy.pp.neighbors” with the first 10 PCs and the size of local neighbourhood specified as 30. Unsupervised clustering of cells was performed using “scanpy.tl.louvain” function with resolution 1.0. Finally, coarse-grained connectivity structures connecting the computed clusters of cells was mapped using “scanpy.tl.paga” with default parameters.

#### scVelo

RNA velocity analysis was performed using scVelo version 0.1.16 (33). Only day 2-day 4 cells from mGVHD2 dataset were included in the analysis. All parameters were kept at default, except: 3,000 genes, 20 PCs, and 30 neighbours considered for RNA velocity estimation. Calculated velocity was projected onto pre-computed bGPLVM embeddings.

#### Gene signature scoring

The “AddModuleScore” function from Seurat was used to calculate gene signatures. The cell cycle score was calculated using 226 cell cycle genes derived from Cyclebase (34), the aerobic glycolysis score used 41 genes associated with the Gene Ontology (GO) I.D. GO:0006096 and the oxidative phosphorylation score used 30 genes associated with I.D. GO:0006119.

#### Gene expression imputation

Gene expression inference from missing data, or imputation, was performed using ALRA (initial release) (https://github.com/KlugerLab/ALRA/)(19) and MAGIC v1.5.0 (35).

#### Differential gene expression analysis and Gene ontology term enrichment analysis

Differential gene expression analysis (DGEA) was performed using “FindMarkers” function from Seurat at default parameters. Comparisons were done as follows: (1) trajectory II cells and trajectory III cells, (2) day 0 and day 1 cells, and (3) day 1 and day 2 cells. (1) was performed using mGVHD2 dataset as input, and (2) and (3) were performed using the integrated mGVHD2 and mGVHD3 datasets as input. Genes from (2) and (3) with Bonferroni adjusted P-value below 0.01 and average log fold change greater than 0.5 were considered as input for gene ontology (GO) term enrichment analysis. GO terms were obtained from “org.Mm.eg.db” Bioconductor annotation package 99 (36). Fisher’s exact test was performed using “goana” function from edgeR (37) to identify over-represented GO terms. Input genes for DGEA was used as the background gene set.

#### Other packages

scRNA-seq data was primarily visualised with the ggplot2 v3.2.1 R package (38). Additional functions were provided by cowplot v0.9.2, in particular via Seurat’s inbuilt plotting features (31). 3D projections were created by the scatterplot3d v0.3-41 R package (39). 3D rotation of projections generated using https://codepen.io/etpinard/pen/mBVVyE?editors=0010. Seurat v3.0.1 was used for conversion of data to loom format.

#### Study approval

All animal procedures and protocols were approved (A0412-617M; P832) and monitored by the QIMR Berghofer Medical Research Institute Animal Ethics Committee.

## Author Contributions

JAE designed research studies, conducted wet-lab experiments, acquired data and analysed data. HJL and CGW designed research studies and performed bioinformatics analyses of data. RK, SO, LIML, MSFS, SBA, AV, MK, AH designed research studies and conducted experiments. VS provided input to the bGPLVM computational modelling of the data. JEP, SAT, GRH, AV and MK provided valuable expertise to the project. SAT, MK and AH conceived the project and interpreted results. AH, JAE, HJL and CGW wrote and edited the manuscript and prepared figures. All authors provided input and approved the manuscript. JAE, HJL and CGW are co-first authors of the manuscript, the order of authors was determined by work load.

## Acknowledgments

This work was supported by research grants from the Australian National Health and Medical Research Council: JEP is supported by an Investigator Grant (1175781); AH, SAT and MK were supported by Project and Ideas Grants (1126399 and 1180951). We would like to thank staff from the QIMR Berghofer flow cytometry facility for their expertise and assistance with experiments, and staff from QIMR Berghofer animal facility for animal care and husbandry. We acknowledge Paul Collins at the QIMR Berghofer sequencing facility, and Scott Wood and his team at the QIMR Berghofer High Performance Computing facility for support with scRNA-seq data management. We acknowledge helpful discussions with Dr. Kate Gartlan (QIMR Berghofer) on matters relating to T cell differentiation during alloSCT. We thank Dr. Kylie R. James (Wellcome Sanger Institute) for reading the manuscript and providing helpful comment on issues relating to CD4^+^ T-cell differentiation.

---

**Supplementary Figure 1.**
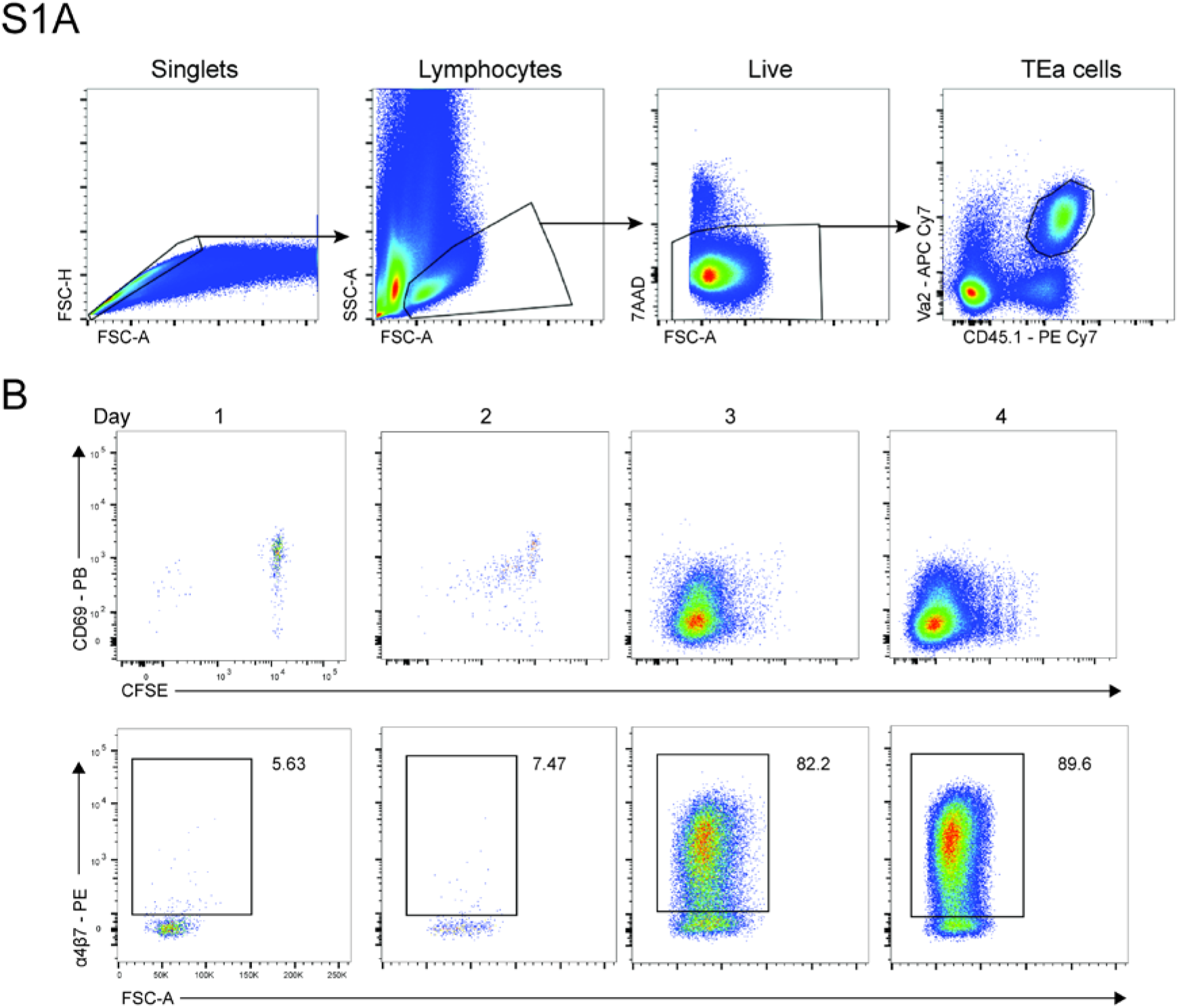
Phenotypic assessment of TEa cells from the mLN. **(A)** FACS gating strategy for isolation of TEa cells (Vα2^+^ CD45.1^+^) from the mLN on day 1- day 4 post-transfer. **(B)** Representative FACS plots showing the expression of CD69, α4β7 and CFSE tracking dye on TEa cells from the mLN on days 1-4 post-transfer.

**Supplementary Figure 2.**
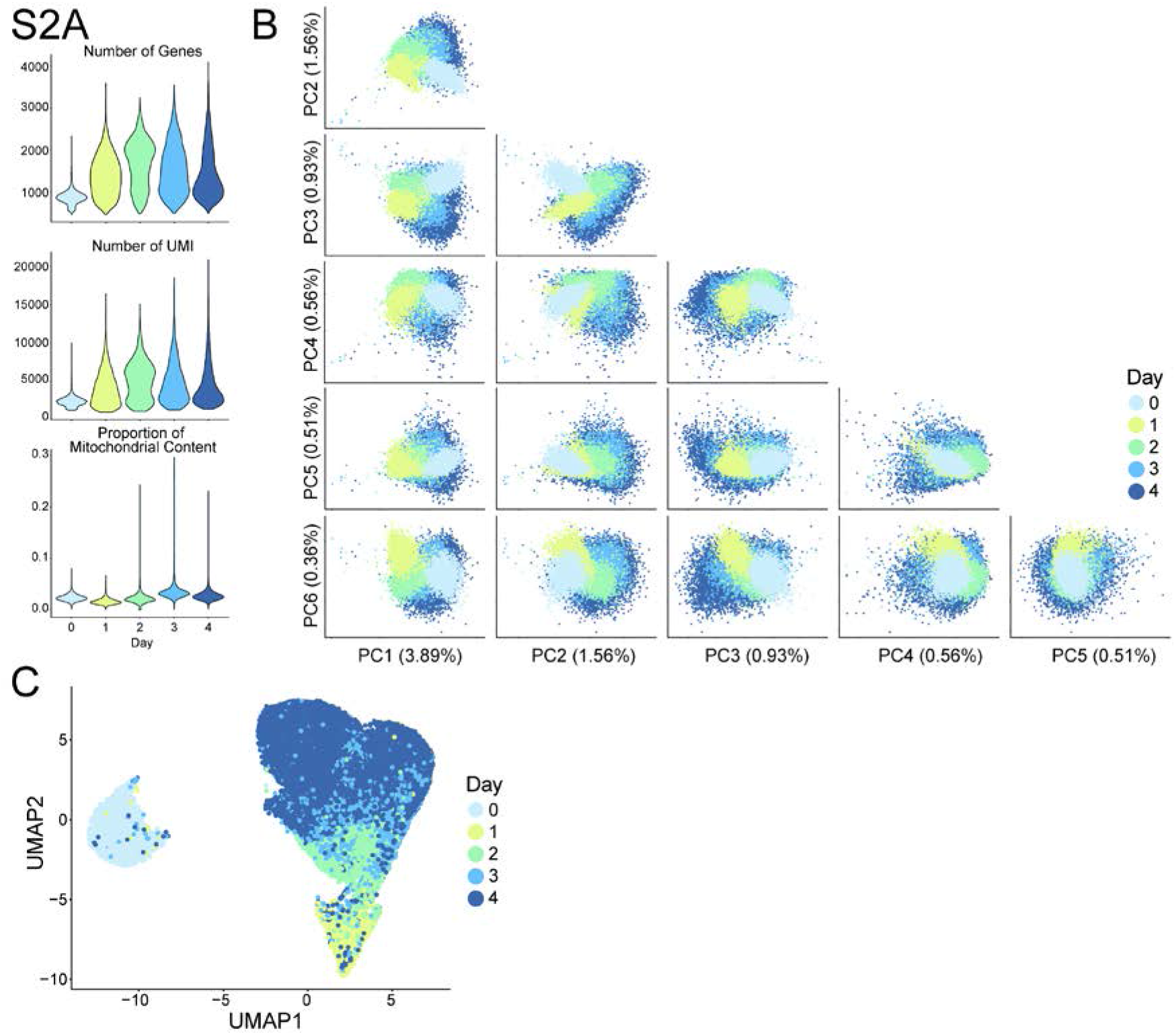
Quality control metrics of scRNA-seq dataset from the mLN. **(A)** Violin plots showing the distribution of TEa cells for number of expressed genes (cells filtered for 200<nGene<6000), number of UMIs and proportion of mitochondrial content (cells filtered for <0.35). **(B)** Principal component analysis (PCA) plots showing TEa cells from the mLN at day 0- day 4 post-transfer. **(C)** UMAP representation of TEa cells from the mLN at day 0- day 4 post-transfer.

**Supplementary Figure 3.**
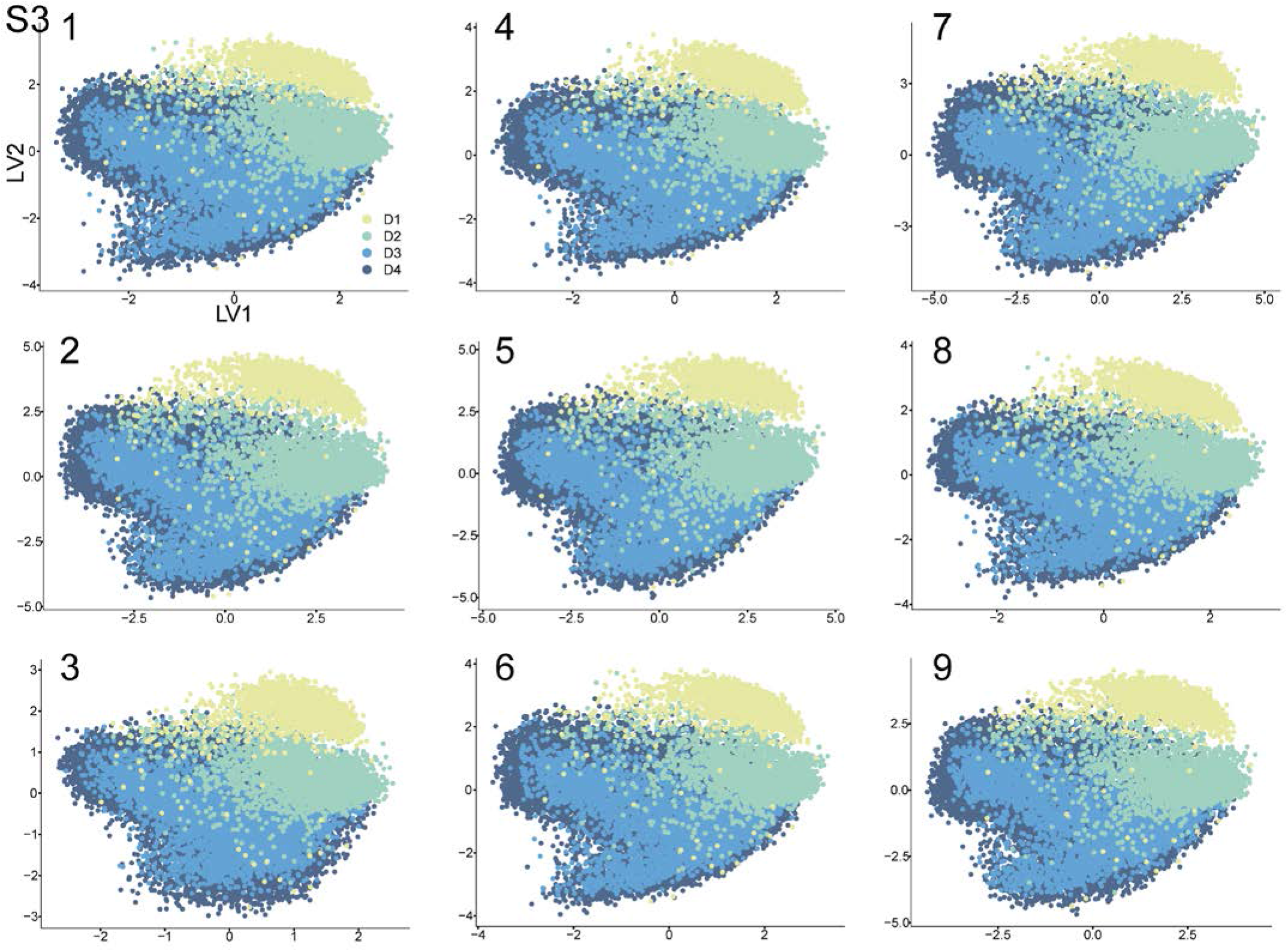
Nine replicates of bGPLVM on the scRNA-seq dataset from the mLN. bGPLVM was run an additional nine times on the day 1-4 post-transfer dataset (mGVHD2). The first two latent variables from each run are displayed. Cells are coloured by timepoint.

**Supplementary Figure 4.**
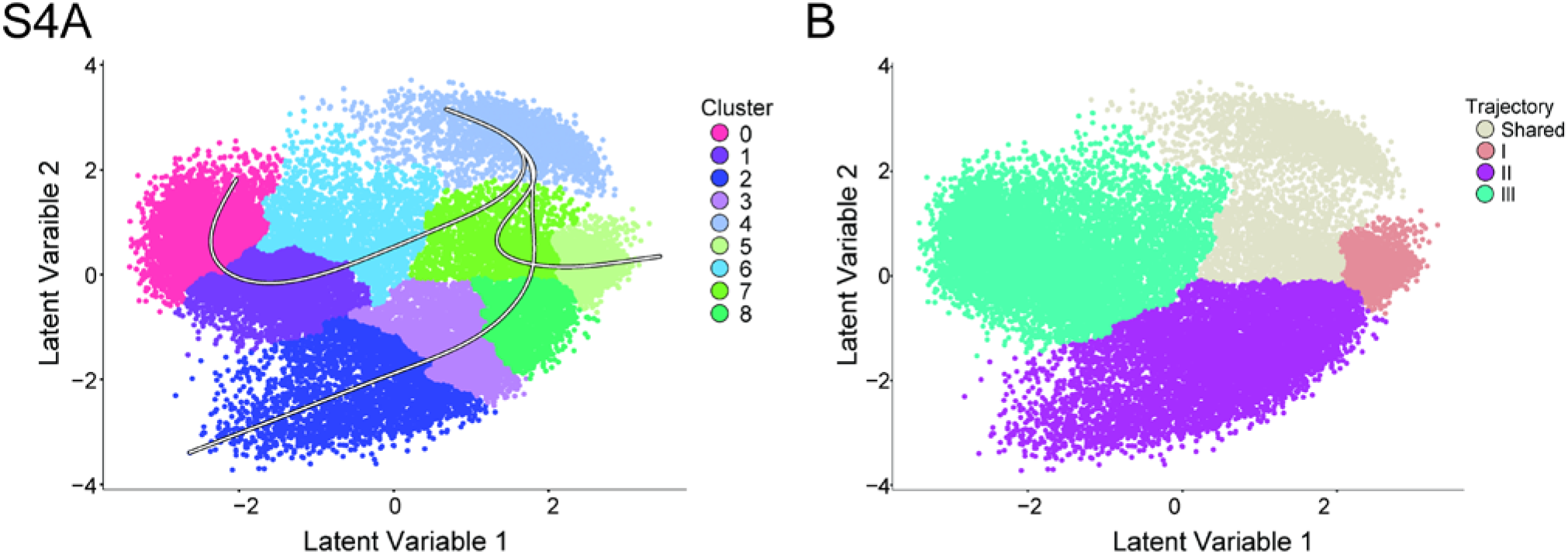
Cluster assignment of the scRNA-seq dataset used for computational modelling with Slingshot. **(A)** Cluster assignment of cells on bGPLVM visualisation used as input for Slingshot to identify the overlaid developmental trajectories. **(B)** bGPLVM representation showing the assignment of cells to either Trajectory I, II, or III derived from Slingshot, or a ‘shared’ cluster which is common to all three trajectories prior to branching.

**Supplementary Figure 5.**
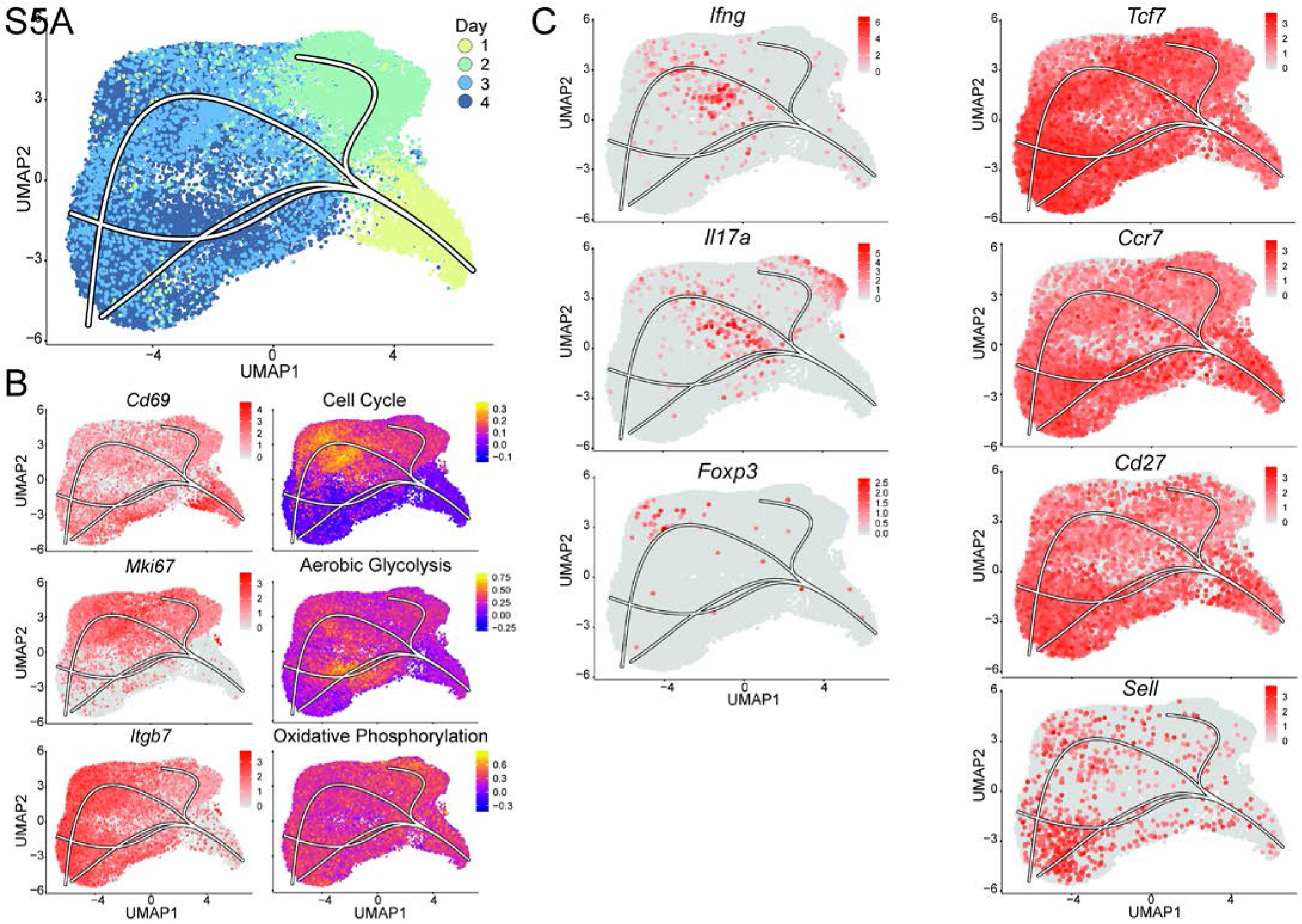
An alternative dimensionality reduction, UMAP, coupled with Slingshot for identification of TEa developmental trajectories. **(A)** UMAP representation of TEa cells on day 1- day 4 overlaid with the developmental trajectories identified by Slingshot. **(B)** Expression of *Cd69*, *Mki67* and *Itgb7* or the cell cycle, aerobic glycolysis and oxidative phosphorylation gene signature scores on the UMAP representation overlaid with Slingshot trajectories. **(C)** Expression of *Ifng*, *Il17a*, *Foxp3*, *Tcf7*, *Ccr7*, *Cd27* and *Sell* on UMAP representation overlaid with Slingshot trajectories.

**Supplementary Figure 6.**
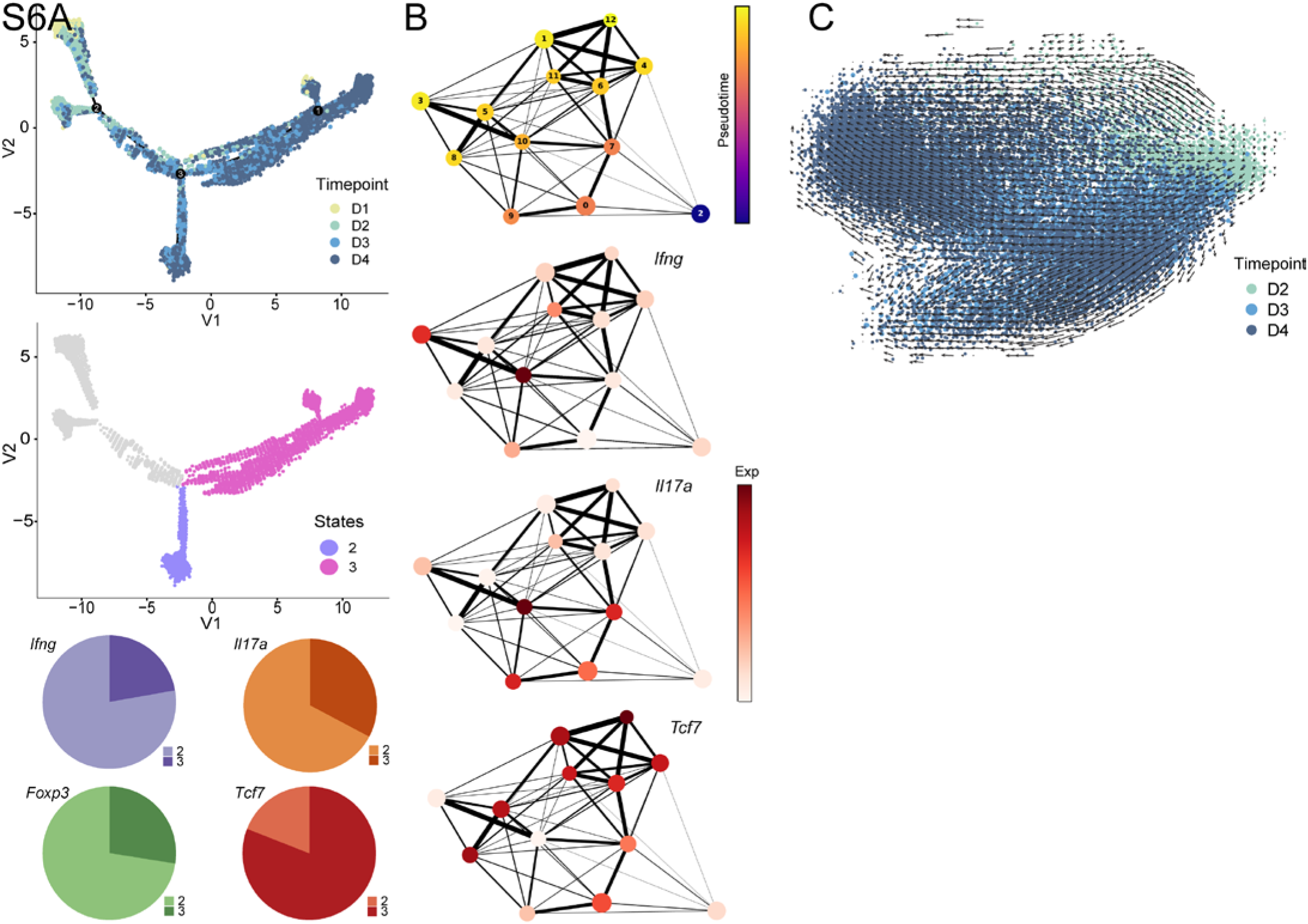
Application of other trajectory inference methods resolves similar developmental bifurcation. **(A)** Monocle DDRTree output with cells coloured by timepoint (*top*) and developmental states of interest (numbered) as their corresponding slingshot trajectory (*middle*). Pie charts (*bottom*) show the number of cells expressing *Ifng*, *Il17a*, *Foxp3*, and *Tcf7* within the states. **(B)** PAGA graph output where each node represents a group of cells characterized by unsupervised clustering and edge weights quantify the connectivity between groups. (*Top*) Cells are shaded by pseudotime and each cluster labelled with a number. Cluster 2 approximately corresponds with day 1 cells. (*Bottom*) Average *Ifng*, *Il17a*, and *Tcf7* expression in each cluster. **(C)** Output from scVelo superimposed on bGPLVM latent variables 1 and 2. Cells shaded by timepoint. Arrows indicate inferred RNA velocity direction and arrow length corresponds to the magnitude of the calculated velocity.

**Supplementary Figure 7.**
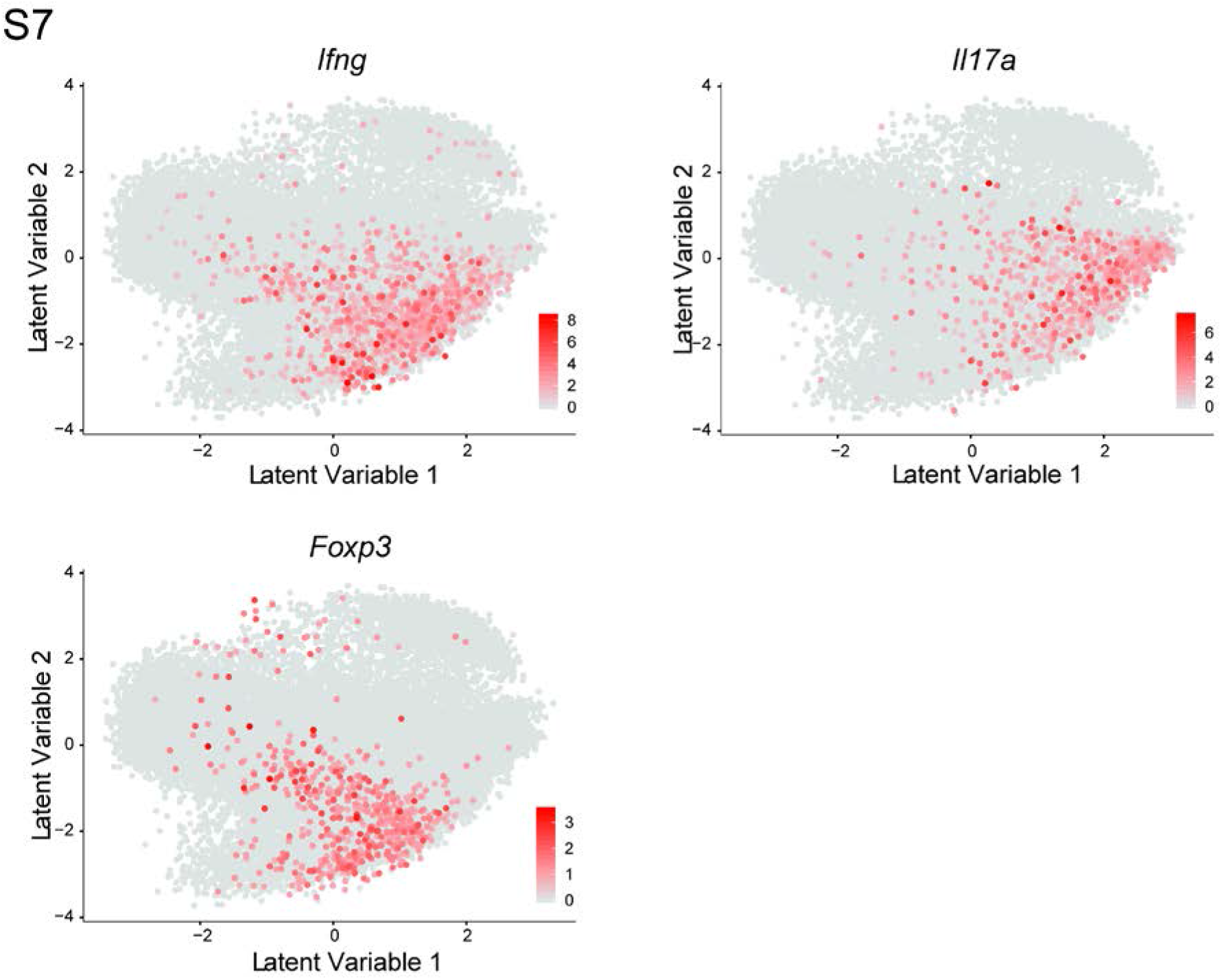
Imputation of cytokine expression does not reveal distinct trajectories among *Tcf7*^lo^ TEa cells. Imputed expression of *Ifng*, *Il17a*, and *Foxp3* by ALRA on bGPLVM latent variables 1 and 2.

**Supplementary Figure 8.**
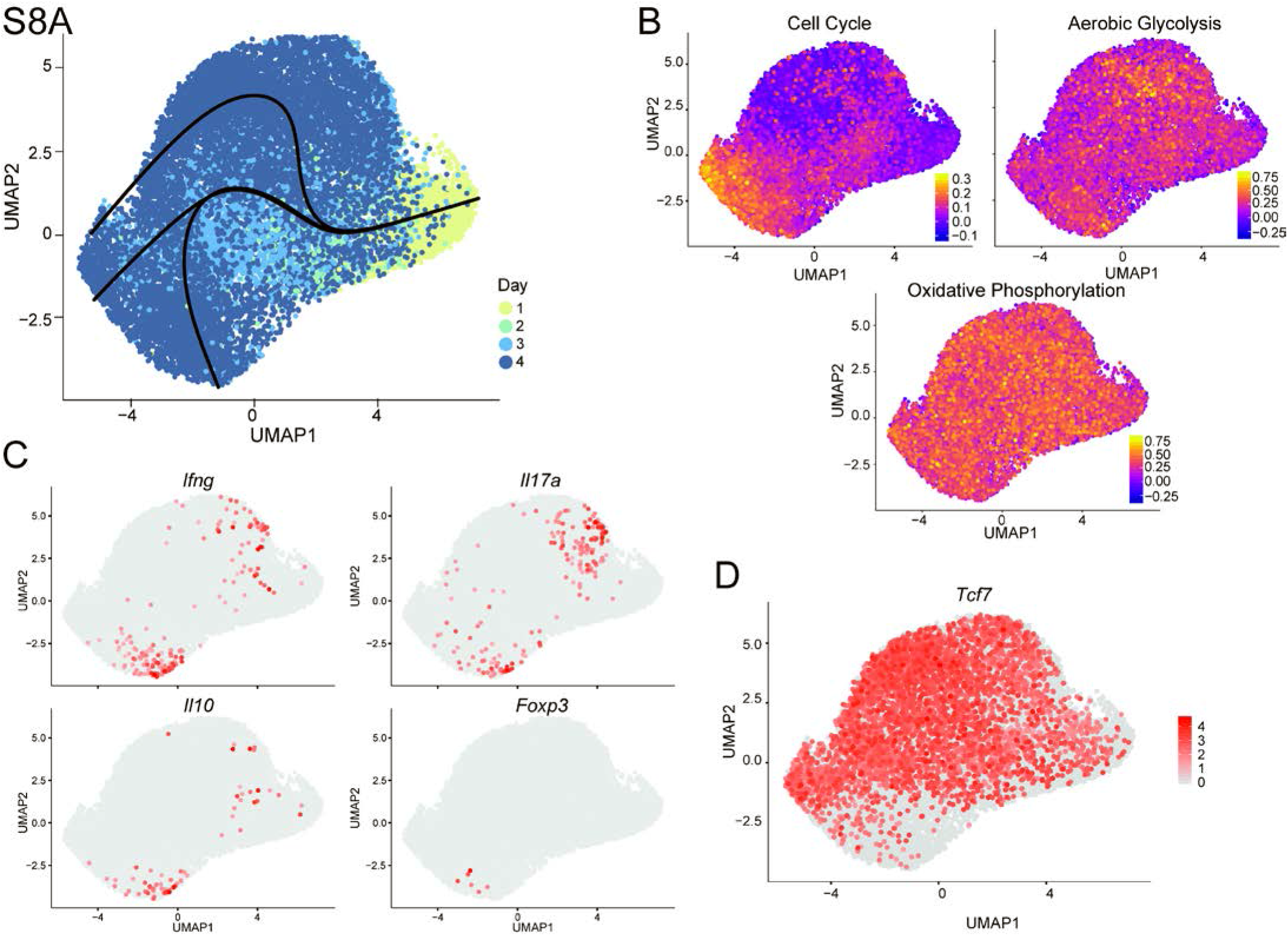
Independent replicate of scRNA-seq from the mLN with UMAP and Slingshot. **(A)**UMAP representation of day 1- day 4 cells in an independent replicate (mGVHD1). Superimposed trajectories generated via Slingshot. **(B)** UMAP representation of day 1- day 4 cells in an independent replicate with *Ifng*, *Il17a*, *Il10*, and *Foxp3* expression superimposed.

**Supplementary Figure 9.**
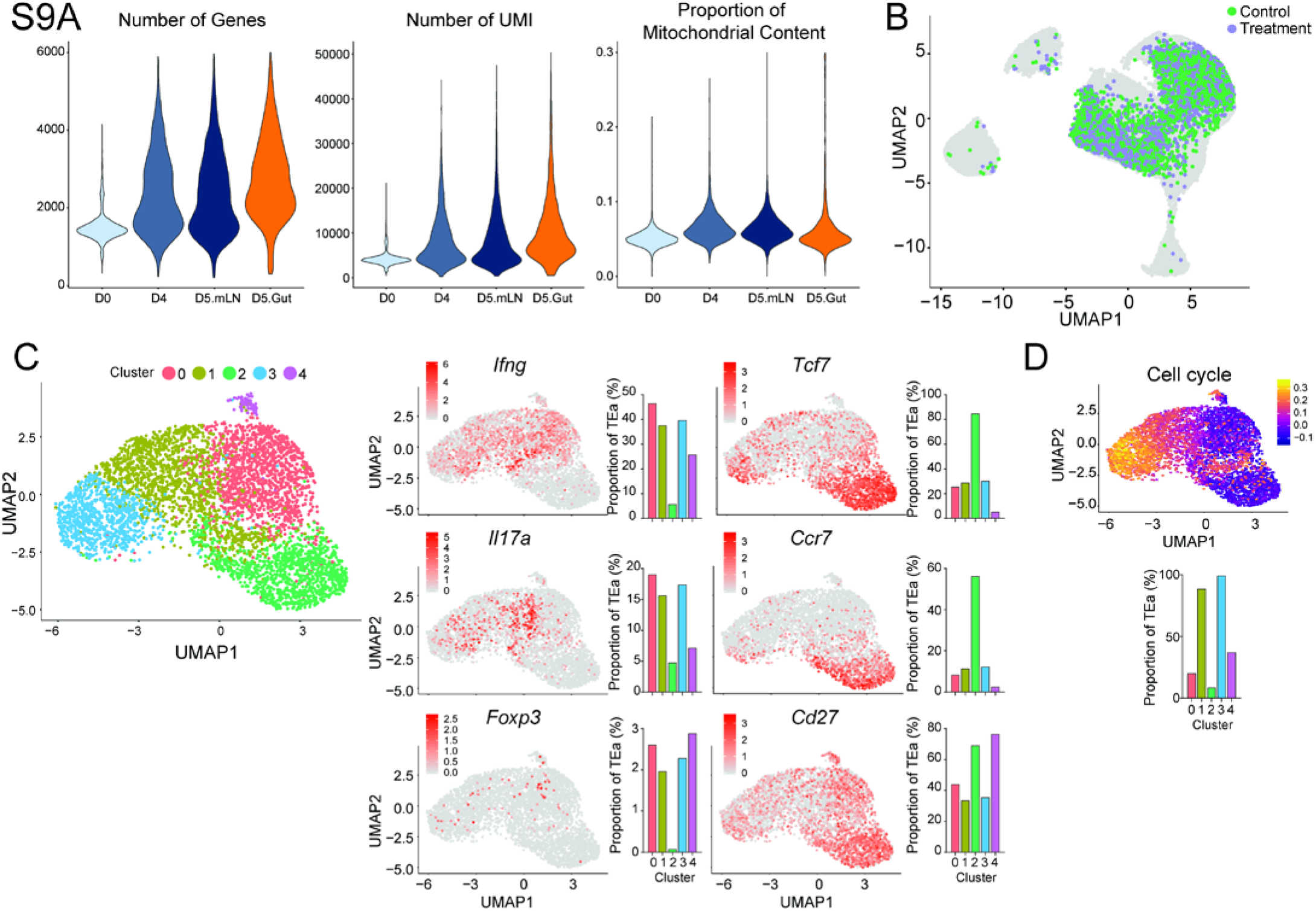
Quality control and preliminary analysis of scRNA-seq dataset from the mLN and gut. **(A)** Violin plots showing the distribution of TEa cells for number of expressed genes (cells filtered for 200<nGene<6000), number of UMIs and proportion of mitochondrial content (cells filtered for <0.3 or <0.15). **(B)** UMAP representation of integrated replicates of scRNA-seq datasets generated using PCA (Seurat) without batch correction. Day 5 post-transplant mLN TEa cells are shaded according to whether they underwent the standard extraction protocol (control) or the extraction protocol for IEL TEa (treatment). **(C)** UMAP representation using PCA (Seurat) of day 5 post-transplant IEL TEa cells, shaded by cluster from unsupervised clustering (Seurat) (*left*). Expression of *Ifng*, *Il17a*, *Foxp3*, *Tcf7*, *Ccr7*, and *Cd27* on UMAP representation (*right*). Bar graphs represent the proportion of cells within each cluster that express each gene. **(D)** Same dataset and embeddings as (C) with the cell cycle gene signature score.

**Supplementary Figure 10.**
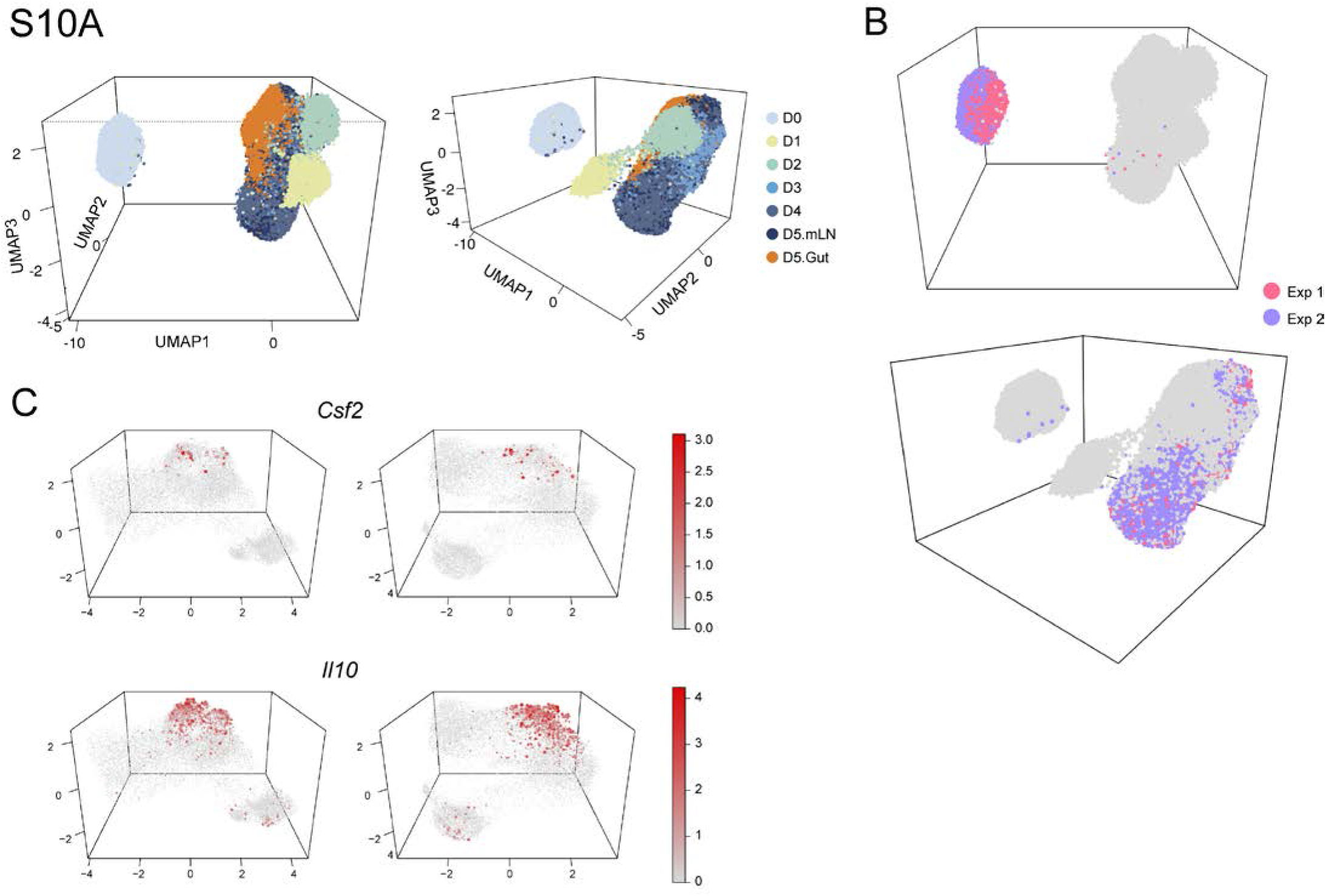
Integration of mLN and gut scRNA-seq experiments. **(A)** Two angles of 3-dimensional UMAP representation produced from 30 scVI latent variables of day 0 – day 5 TEa cells. **(B)** Same UMAP representation as (A) with cells shaded by their experiment of origin, highlighting cells from day 0 (*top*) or from day 4 (*bottom*). **(C)** Two angles of 3-dimensional UMAP representation showing TEa cells from day 2 mLN, day 5 mLN, and day 5 gut, with expression of *Csf2* and *Il10*.

**Supplementary Figure 11.**
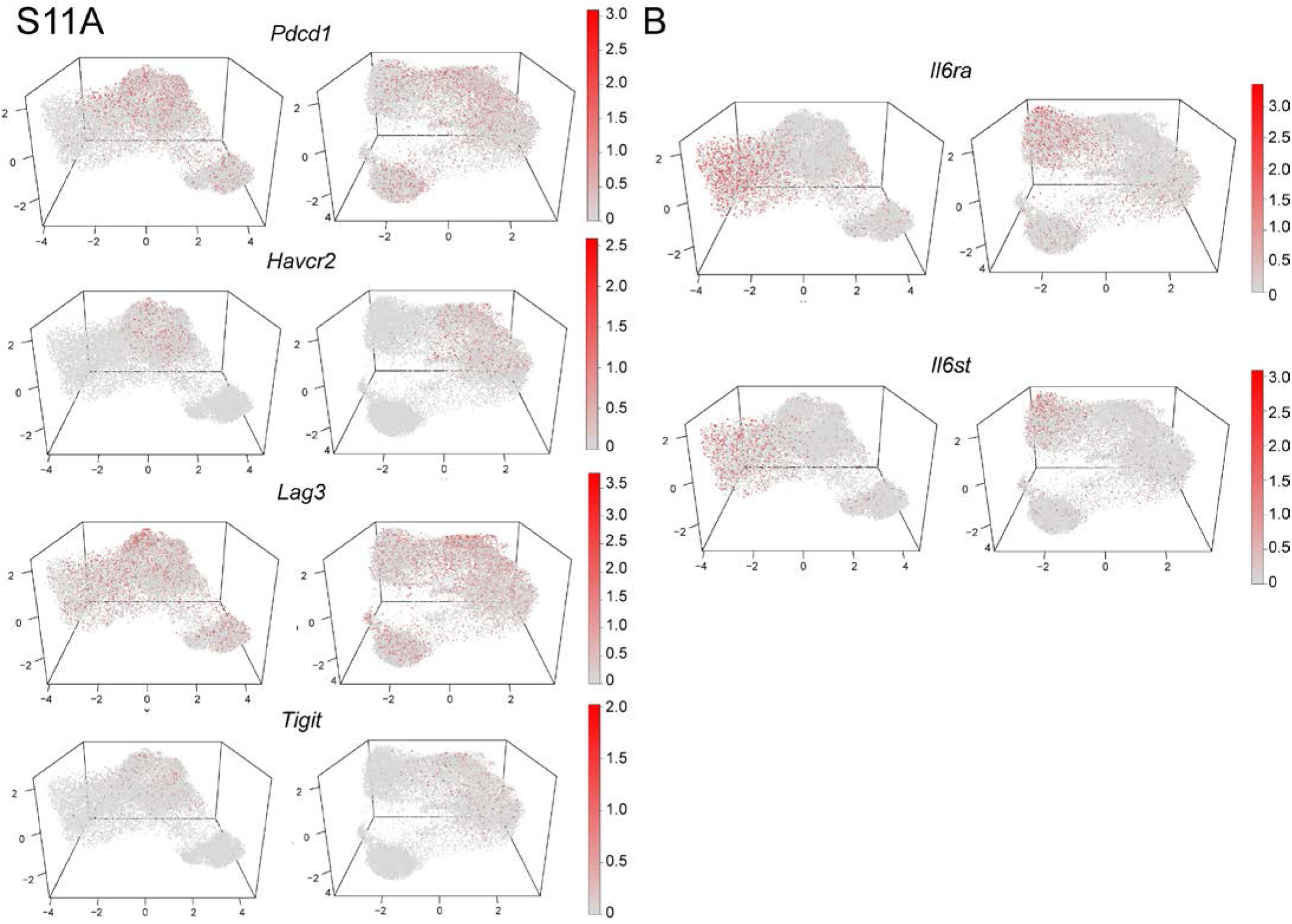
Analysis of expression of genes associated with T cell exhaustion and IL-6 signalling. **(A)** Expression of *Pdcd1*, *Havcr2*, *Lag3* and *Tigit* on 3-Dimensional UMAP visualisation. **(B)** Expression of *Il6ra* and *Il6st* on 3-Dimensional UMAP visualisation.

**Supplementary Figure 12.**
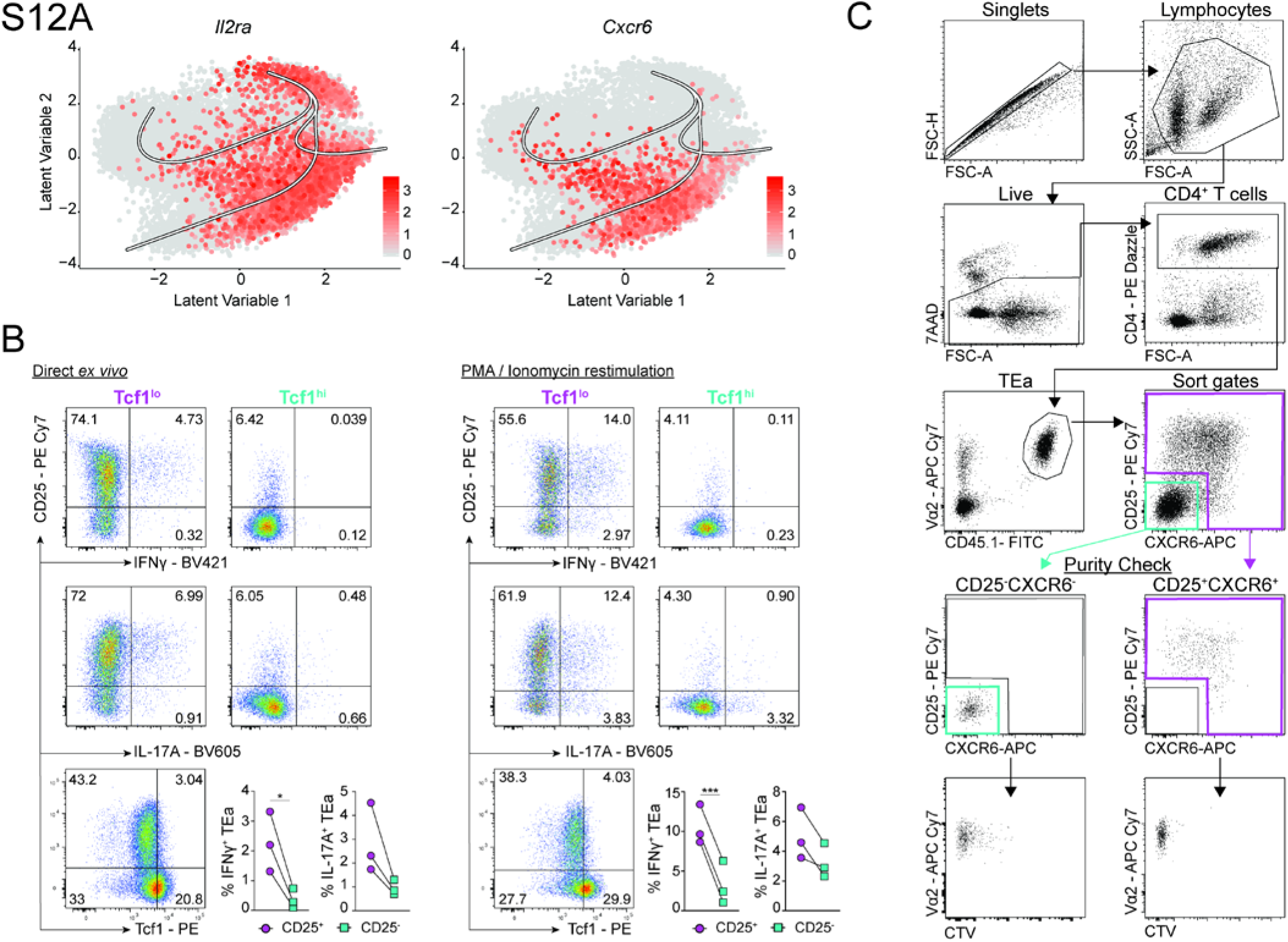
Phenotypic assessment of Tcf1^hi^ and Tcf1^lo^ cells by flow cytometry. **(A)** bGPLVM representation of TEa cells from the mLN from day 1- day 4 showing the expression of *Il2ra* and *Cxcr6*, overlaid with Slingshot trajectories. **(B)** Representative flow cytometry plots showing the expression of CD25, IFNγ, IL-17A and TCF1 on Tcf1^lo^ or Tcf1^hi^ TEa cells from the mLN at day 4 post-transfer, directly *ex vivo* or after restimulation with PMA/ionomycin. Graphs show the percentage of CD25^+^ or CD25^−^ TEa cells that express IFNγ and IL-17A. Data shown are from one experiment (*n* = 3 mice). Statistical analysis was performed using a Paired *t* test. *p<0.05, **p≤0.01, ***p≤0.001, ****p≤0.0001. **(C)** Flow cytometry gating strategy used to isolate CD25^−^CXCR6^−^ and CD25^+^CXCR6^+^ TEa cells from the mLN at day 4 post-transfer. The corresponding purity checks and CTV expression for each population are shown.

